# Mechanistic Insights into the *Plasmodium* Phosphatase UIS2 in eIF2α Dephosphorylation via Integrated Structure-Based Modeling

**DOI:** 10.1101/2025.05.14.654012

**Authors:** Su Wu, Gerhard Wagner

## Abstract

The development of novel antimalarial drugs requires the identification of parasite-specific vulnerabilities. Translation control, regulated by eIF2α dephosphorylation, is essential for *Plasmodium* development during infection. The parasite phosphatase UIS2 regulates this modification. However, the structural basis for UIS2 substrate recognition remains unknown. Here, we show that UIS2 is crucial for parasite development in both liver and blood stages. Its N-terminal and phosphatase domains each independently bind the eIF2α phosphoserine-59 loop through electrostatic interactions. Integrated structural modeling, using AlphaFold and molecular dynamics simulations, reveals a defined binding pocket within the phosphatase domain, with a geometry distinct from human PP1α. This pocket contains six cooperative binding patches that use an electrostatic network to stabilize the phosphopeptide near the active site and contribute to substrate specificity. Metadynamics simulations show that the inhibitor Salubrinal competes with the phosphopeptide for binding to this pocket. Molecular docking and free energy perturbation analyses show that Salubrinal targets three of these binding patches through hydrophobic and hydrogen-bonding interactions, blocking substrate binding. These findings show the structural and energetic basis of eIF2α dephosphorylation by UIS2. They provide an integrated modeling approach for phosphatase-substrate interactions and a mechanistic rationale for selective inhibition of a parasite-specific translation regulator.

## Introduction

Malaria, caused by *Plasmodium* parasites, is a major global health burden, with 263 million cases and 608,000 deaths reported in 2024.^1^ During transmission, infected mosquitoes inject sporozoites into the bloodstream, which migrate to the liver, replicate, and release merozoites back into circulation (Fig. S1A).^2^ Merozoites then invade red blood cells, progress through ring, trophozoite, and schizont stages, and rupture the host cell to release new parasites, triggering inflammation and anemia. Artemisinin-based combination therapies (ACTs) are the primary treatment, but artemisinin-resistant *Plasmodium* strains harboring *Kelch13* mutations, which impair drug activation, are now widespread.^3,4^ This growing resistance highlights the urgent need to identify key regulators of parasite growth. Translation initiation factors, critical for protein synthesis across the parasite life cycle, are promising therapeutic targets.^5^

Up-regulated in Infective Sporozoites 2 (UIS2)^6^ is a phosphatase that dephosphorylates eukaryotic initiation factor 2α (eIF2α) to enable translation initiation. Functional studies show that *UIS2*-null sporozoites can invade hepatocytes but fail to develop into hepatic-stage parasites, thereby blocking further progression of infection.^7^ UIS2 localizes to the parasitophorous vacuole membrane (PVM) in both hepatic and blood stages via its N-terminal RILDE motif, which depends on additional proteins for anchoring.^8-10^ By dephosphorylating eIF2α, UIS2 allows eIF2B to convert GDP-bound eIF2 to its active GTP-bound form (Fig. S1B). Active eIF2-GTP loads the initiator methionyl-tRNA (Met-tRNA_i_) onto the 40S ribosomal subunit, forming the 43S pre-initiation complex that scans mRNA for the initiation codon. eIF2α phosphorylation at serine 59 (P-eIF2α) by kinases eIK1, eIK2, or PK4 under various stress conditions inhibits eIF2B,^11-14^ preventing eIF2 recycling and translation.^15^ Although eIF2α dephosphorylation is essential during the parasite life cycle, UIS2’s role in this process during the symptomatic blood stage remains poorly understood.

The purified UIS2 phosphatase domain (PD) displays Mn^2^+-dependent enzymatic activity and dephosphorylates P-eIF2α *in vitro* without cofactors, demonstrating intrinsic substrate specificity. UIS2’s N-terminal domain (NTD) binds P-eIF2α but not non-phosphorylated eIF2α, independently of the PD, suggesting a two-step recognition mechanism.^7^ Salubrinal,^16^ an inhibitor, blocks UIS2-mediated eIF2α dephosphorylation *in vitro* and *in vivo* and, when combined with artemisinin, suppresses parasite recrudescence in animal models.^17^ Despite these findings, key questions remain: what is the structural basis of UIS2’s substrate recognition and catalysis? How does the PD bind the flexible pSer59 loop to achieve specificity? Does Salubrinal directly interfere with this interaction? However, experimental capture of the UIS2–eIF2α complex is challenging because serine/threonine phosphatases process and release substrates within milliseconds (kcat ≈ 1–100 s^−1^), making the complex too short-lived for crystallography or routine cryo-EM.^18^ For example, the active site structure of PP1α, the eIF2α phosphatase in humans, was solved by crystallography only after stabilization with the catalytic inhibitor tautomycetin.^19^ Cryo-EM of PP1α-GADD34-eIF2α-actin complex reveals substrate binding but lacks resolution to define specific interactions.^20^ To bypass these experimental limitations, we applied physics-based computational modeling to investigate UIS2 structure, conformational dynamics, and substrate interaction.

In this study, we analyzed genome-wide loss-of-function screen data and single-cell transcriptomes across the *Plasmodium* life cycle. We show that UIS2 is essential for blood-stage growth and is highly expressed in both hepatic and erythrocytic forms. AlphaFold (AF) models of UIS2 predict a purple acid phosphatase-like fold with a defined anionic cavity and conserved active site residues. AF predicts two UIS2–eIF2α complexes, where either the NTD or PD binds the Ser59 loop via electrostatic interactions. AF models, followed by molecular dynamics (MD) simulations, refine phosphopeptide orientation and binding at the UIS2-PD interface. The UIS2-PD substrate-binding pocket adopts a geometry distinct from that of PP1α, stabilized by an extensive network of hydrogen bonds and ionic interactions that define substrate specificity. Trajectory analysis identifies six cooperative patches within the UIS2-PD pocket that secure substrate binding. Metadynamics and free-energy perturbation (FEP) calculations show that Salubrinal displaces the phosphopeptide by binding to three of these patches via hydrophobic and hydrogen-bonding interactions. These findings provide a structural and energetic framework for UIS2’s substrate specificity and offer insights for designing inhibitors targeting *Plasmodium* translation control.

## Results

### UIS2 is essential for *Plasmodium* proliferation and is highly expressed in blood and liver *stages*

We investigated translation initiation factors in *Plasmodium berghei*, a rodent malaria parasite, with symptoms resembling those of *Plasmodium falciparum*, the primary human malaria parasite. We analyzed *in vivo* phenotype screening data from PlasmoGEM, which assessed the growth of *P. berghei* mutants, each mutant lacking one of 2,578 *Plasmodium* genes, during erythrocytic stages in mice.^21^ *UIS2* and translation initiation factors *eIF4G, eIF4A3, eIF3D*, and *eIF3I*, are essential for *P. berghei* growth (Fig. 1A). Previous studies showed that *UIS2* disruption in *P. falciparum* led to parasite death in cultured human erythrocytes.^10^ Knockout of eIF2α kinases *eIK2* and *PK4* resulted in slow growth (Fig. 1A), indicating they are dispensable during erythrocytic stages. These findings confirm that *UIS2* is critical for parasite proliferation in the erythrocytic stage, beyond its known role in sporozoite-to-liver transition.

**Fig. 1:**
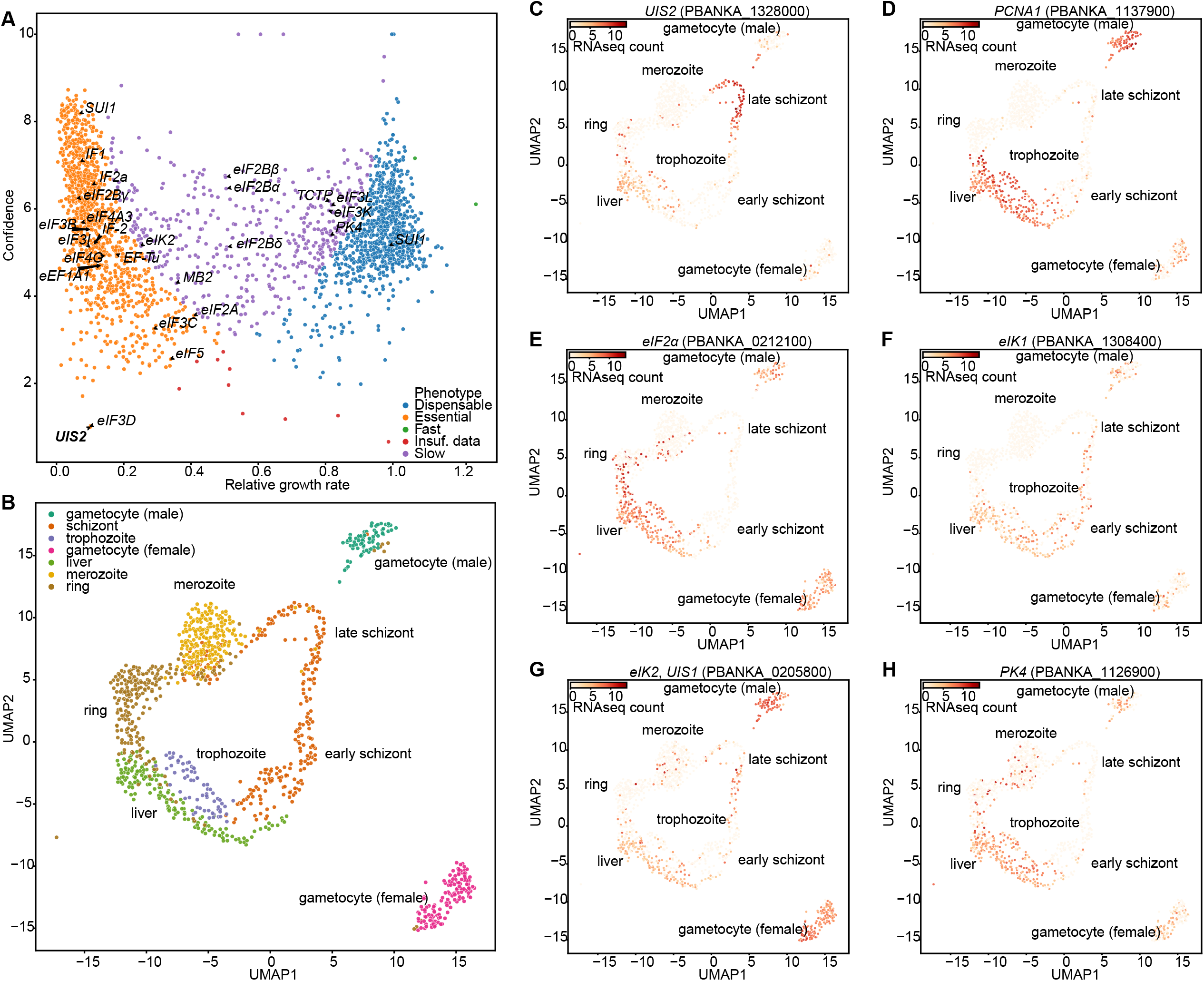
*UIS2* is critical for *Plasmodium* growth in the erythrocytic stage and is highly expressed in erythrocytic and hepatic stages. **(A)** The dot plot shows relative growth rates of 2,578 *Plasmodium* berghei mutants, each with a single gene knocked out, following pooled infection in mice with schizont-stage parasites. The x-axis displays growth rates, calculated from changes in barcoded knockout gene abundance, normalized to the control post-infection, using next-generation sequencing. The y-axis shows confidence, expressed as the negative logarithm of variance, derived from barcode counts across three replicate infections. Genes are color-coded by phenotype after knockout: essential, slow-growing, dispensable, or fast-growing. Translation initiation factors, including UIS2, are annotated. **(B)** UMAP projection of 1,137 single *P. berghei* parasites spanning all life cycle stages in the mouse host, clustered by transcriptomic similarity based on single-cell RNA-seq. Each dot represents one parasite, colored by developmental stage. **(C–H)** UMAP plots show expression of *UIS2* and related genes across parasite life stages, based on clusters defined in **(B)**. Parasites with detectable expression of each gene are shown in red, with color intensity reflecting transcript abundance from single-cell RNA-seq data.

We examined *UIS2* mRNA expression data across the parasite life cycle from the Malaria Cell Atlas,^22^ which provides single-cell transcriptomic profiles of *P. berghei* parasites across morphological stages in mice (Fig. 1B). *UIS2* expression is elevated in the erythrocytic ring form, trophozoite and hepatic stages (Figs. 1C). These stages are critical for parasite proliferation, as shown by co-expression of DNA replication markers *PCNA1* and *ORC1* (Figs. 1D and S2A). High *UIS2* expression also occurs in late schizonts, where the parasite completes nuclear division and prepares for egress, alongside markers *CDPK5* and *HSP101* (Figs. S2B, C).^23,24^ In sporozoites within mosquito salivary glands, *UIS2* is transcriptionally active but repressed at the translational level by the RNA-binding protein Puf2, until transmission to the host.^7^ We observed strong co-expression of *Puf2* with *UIS2* in mosquito sporozoites but low *Puf2* expression in mouse parasite stages (Figs. S3A-D). These results indicate that UIS2 supports hepatic and erythrocytic parasite development through transcriptional upregulation and stage-specific relief from Puf2-mediated repression.

We also investigated mRNA expression of *eIF2α* and its kinases *eIK1, eIK2*, and *PK1*. Their transcripts are elevated in hepatic stage and erythrocytic trophozoites (Figs. 2E-H). In erythrocytic stages, mRNA levels typically reflect protein expression, unlike in sporozoites where transcription and translation are often uncoupled.^25^ Thus, elevated transcripts of *UIS2, eIF2α*, and its kinases suggest increased protein levels and coordinated regulation of eIF2α phosphorylation during erythrocytic development. High expression of translation initiation factors, such as *eIF4E, eIF4G*, and *eIF3D*, in hepatic stage and erythrocytic trophozoites (Figs. S2D-F), further supports heightened translation initiation activity during parasite proliferation. These findings show that UIS2 controls eIF2α phosphorylation in hepatic parasites, erythrocytic trophozoites, and late schizonts, contributing to malaria symptoms.

**Fig. 2:**
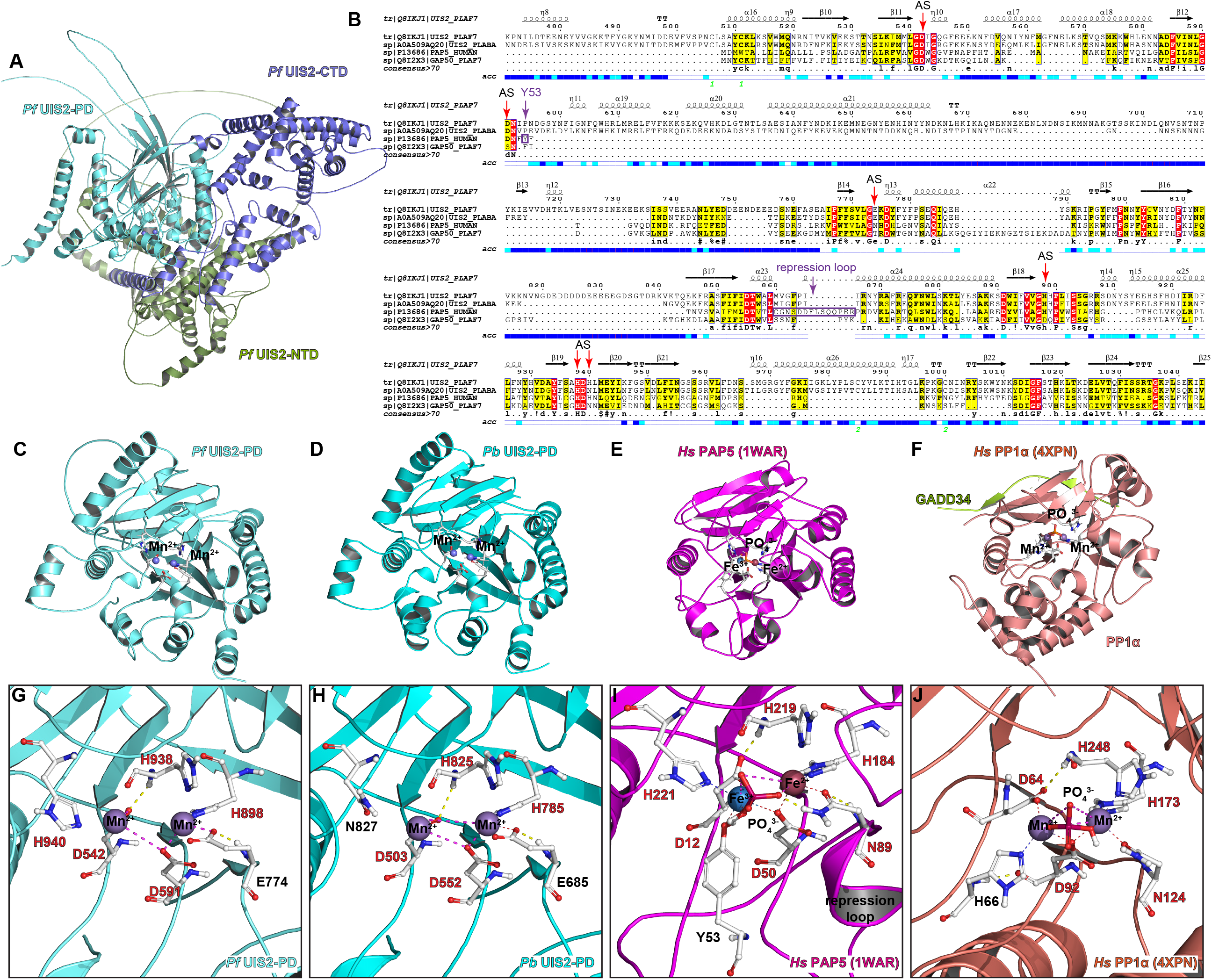
The phosphatase domain of UIS2 shows both sequence and structural homology with purple acid phosphatase. **(A)** Predicted protein structure of full-length *Pf* UIS2 by Alphafold3. **(B)** Multiple sequence alignment of UIS2-PDs from *P. falciparum* and *P. berghei*, human PAP5, and *P. falciparum* GAP50. The alignment was performed using Clustal Omega and Escript 3.0. Secondary structure elements are indicated: a-helices (squiggles), 3_10_-helices (small squiggles), b-strands (arrows), b-turns (TT letters). Identical residues are highlighted in red boxes, and similar but significantly different residues are highlighted in yellow boxes. Solvent accessibility is indicated below the alignment: blue (exposed), cyan (partially exposed), and white (buried). Green numbers mark predicted disulfide bridges. Active-site residues are labeled “AS” and marked with red arrows. Tyr53 and the repression loop of human PAP5 are highlighted in purple. **(C-F)** Cartoon representation of *Pf* UIS2-PD, *Pb* UIS2-PD, *Hs* PAP5 and *Hs* PP1α protein structures. The phosphate ions are shown as orange sticks. **(G-J)** Close-up views of the active site among *Pf* UIS2-PD, Pb UIS2-PD, *Hs* PAP5 and *Hs* PP1α. Conserved catalytic residues shared among all four phosphatases are highlighted in red.

### UIS2 contains a purple acid phosphatase–like fold in its phosphatase domain

To understand how *Plasmodium* UIS2 mediates eIF2α dephosphorylation, we used AF to model the full-length UIS2 proteins from *P. falciparum* (*Pf* UIS2) and *P. berghei* (*Pb* UIS2). Both proteins comprise three domains: NTD, a central PD, and a C-terminal helical domain (CTD) (Fig. 2A; Figs. S4A, B, E, F). *Pf* and *Pb* UIS2-PDs were predicted with high local confidence (pLDDT > 90), indicating a well-folded enzymatic domain (Figs. S4C and S4G). NTDs and CTDs showed lower confidence, suggesting structural flexibility or disorder. High interdomain confidence scores (ipTM = 0.89, *Pf* UIS2; 0.92, *Pb* UIS2) indicated accurate modeling of domain contacts, while moderate global accuracy (pTM = 0.69, *Pf* UIS2; 0.65, *Pb* UIS2) suggested flexibility in domain orientation. Residue–residue distance error plots confirmed that the three domains are separated by flexible linkers (Figs. S4D and S4H).

AF used HHsearch for template search and identified structural homologs of the UIS2-PD protein from the Protein Data Bank (PDB). HHsearch identified human purple acid phosphatase (PAP5; PDB: 1WAR) and *P. falciparum* GAP50 (PDB: 3TGH) as top structural homologs for UIS2-PDs (Fig. 2B). Sequence alignment confirmed conserved motifs among *Pf* and *Pb* UIS2-PDs, PAP5 and GAP50. However, a unique insertion (residues 595–755) was present only in *Plasmodium* UIS2-PDs but not PAP5 or GAP50 (Fig. 2B). PP1α showed lower sequence similarity and was not among the top structure matches.

We next compared the protein structures of *Pf* and *Pb* UIS2-PDs with PAP5 and PP1α. All four contain a central β-sandwich surrounded by α-helices and a surface-accessible concave pocket near the active site (Figs. S5A, C, E, G). Although UIS2-PDs and PAP5 share overall architecture, the surrounding helices differ in orientation (Figs. 2C-E, S5B, D, F). The active site of UIS2-PDs, coordinating two Mn^2^+ ions, is buried within the β-sandwich and accessed via a surface cavity (Fig. S5A and S5C). PAP5 contains a similar catalytic core but coordinates Fe^2+^/Fe^3+^ ions instead (Fig. 2E, S5E, and S5F).^26,27^ PP1α (PDB: 4XPN) contains a central β-sandwich and a catalytic center with two Mn^2^+ ions, but differs from UIS2-PD in secondary structure arrangement and active-site geometry (Figs. 2F, S5G and S5H).

To further assess active-site conservation, we analyzed metal-binding residues in UIS2-PDs (Figs. 2G–J). *Pf* and *Pb* UIS2-PDs contain histidines (His898/938 in *Pf*; His785/825 in *Pb*), aspartic acids (Asp542/591 in *Pf*; Asp503/552 in *Pb*), and additional residues (Glu774 and His940 in *Pf*; Glu685 and Asn827 in *Pb*) that align closely with the active-site residues of PAP5 (His184, His219, His221, Asp12, Asp50, Asn89) (Fig. 2I), indicating conserved catalytic features.^27^ PAP5 has unique features, such as Tyr53, which forms a Fe^3^+–tyrosinate charge-transfer complex that gives rise to the enzyme’s purple color,^28^ and a cleavable regulatory repression loop (residues 140–153) that can block substrate access.^29^ In contrast, PP1α coordinates Mn^2^+ through Asp64, His66, Asp92, His173, Asn124, and His248 (Fig. 2J), and its active-site geometry differs from both PAP5 and UIS2-PDs. GAP50, while structurally similar, lacks essential catalytic residues and does not exhibit phosphatase activity (Figs. S5I–K).^30^ It coordinates Co^2^+ ions differently, with limited conservation of active-site residues (e.g., Asp34, His256) (Fig. S5L). Together, these findings show that UIS2-PD shares overall structural homology with PAP5, coordinates Mn^2^+ similarly to PP1α, and preserves key catalytic features found in both enzymes.

### NTD and PD of UIS2 independently bind the S59-loop of eIF2α via electrostatic interactions

We next investigated how UIS2 recognizes P-eIF2α through structural modeling. Sequences of *Pf* eIF2α and *Pb* eIF2α are highly similar to human eIF2α, with conserved domains including the S1 domain, an α-helical subdomain, and a C-terminal domain (Figs. 3A, S6A). The phosphorylated Ser59 residue is located within a loop region of the S1 domain. AF modeling yielded UIS2 structures with high confidence (pLDDT >70), especially in the S1 domain (pLDDT >90; Figs. S6B–G). These predicted structures resembled the previously reported NMR structure of human eIF2α.^31^

**Fig. 3:**
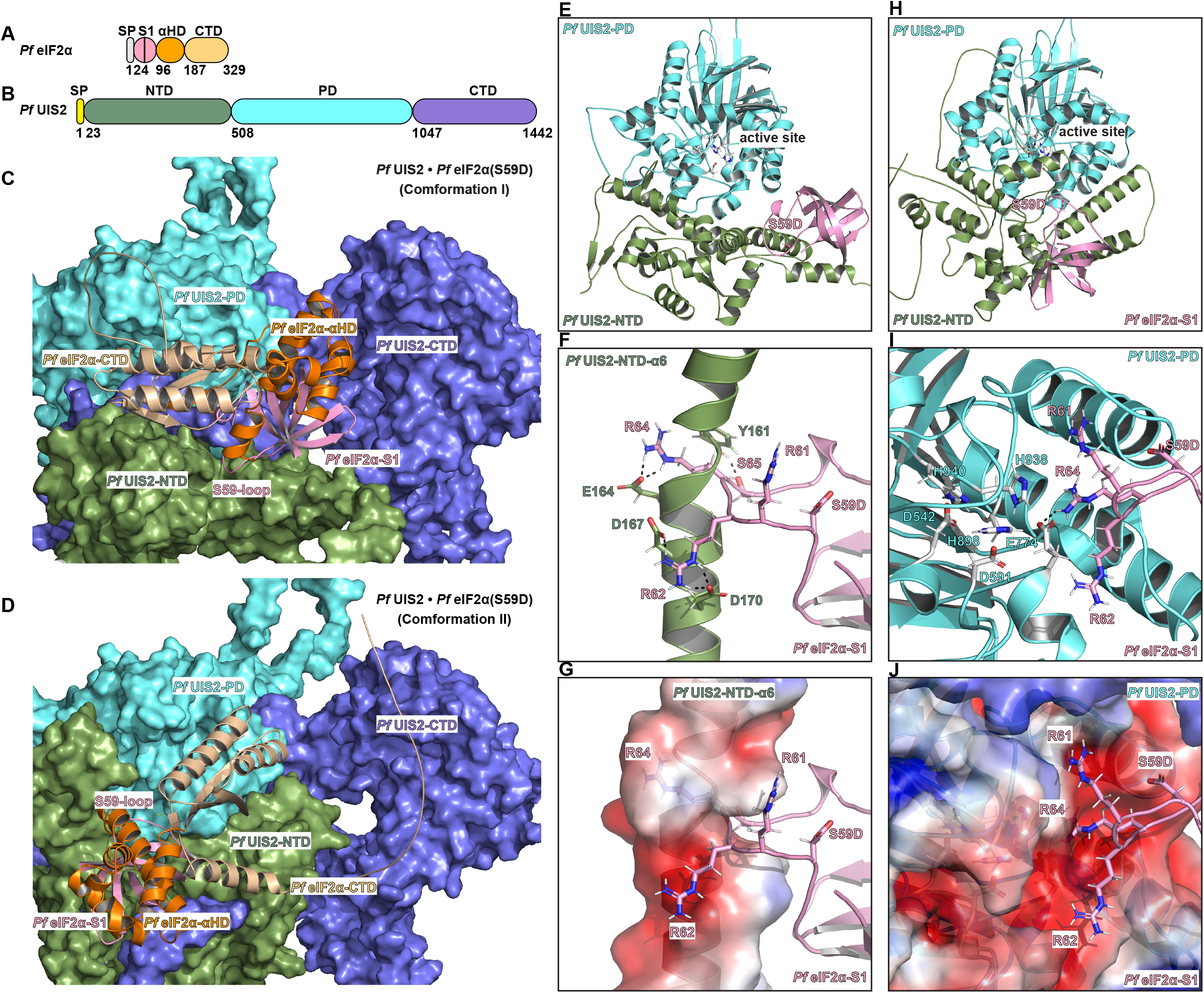
NTD and PD of UIS2 independently bind the S59-loop of eIF2α via electrostatic interactions. **(A)** Domain organization of *Pf* eIF2α. with Ser59 located in the S1 domain (black line). **(B)** Domain organization of the full-length *Pf* UIS2. **(C)** Predicted complex structure (conformation I) of *Pf* UIS2 bound to *Pf* eIF2α S59D (a phosphomimetic mutation), determined by AlphaFold2.3 multimer. **(D)** Predicted complex structure (conformation II) of UIS2 bound to eIF2α S59D. **(E)** Isolated view of conformation I showing UIS2-NTD and UIS2-PD together with the eIF2α S1 domain. Active-site residues of UIS2-PD are shown as silver sticks. **(F)** Close-up of the interface between *Pf* UIS2-NTD helix α6 and the S59-loop of the *Pf* eIF2α S1 domain. Polar contacts are indicated by dashed lines. **(G)** Electrostatic surface map of UIS2-NTD helix α6 (red, negative charges; blue, positive). The eIF2α S59-loop is shown in cartoon, with interacting residues as sticks. **(H-J)** Isolated views of conformation II showing interactions between UIS2-PD and the S59-loop of eIF2α.

To understand the structural basis between UIS2 and P-eIF2α interaction, we used AF2-Multimer to model complexes between a phospho-mimetic mutant (Ser59Asp) of *Pf* eIF2α and *Pf* UIS2 (Fig. 3B). AF2-multimer is optimized for inter-chain interaction prediction,^32^ and it better recapitulated published biochemical data at the UIS2-NTD–eIF2α S1 loop interface.^7^ AF2-multimer generated two distinct top-ranked conformations, each showing the S59-loop of *Pf* eIF2α interacting with either UIS2-NTD or UIS2-PD (Figs. 3C, D). The interaction region displays relatively low pLDDT scores, likely due to structural flexibility in the S59-loop of eIF2α (Figs. S7A-D).

In the first conformation, the NTD and CTD cover the UIS2-PD active site, and the S59-loop interacts directly with the UIS2-NTD (Fig. 3C). This finding is consistent with prior biochemical evidence that UIS2-NTD specifically binds phosphorylated eIF2α.^7^ The interaction involves the α6 helix of UIS2-NTD, located distally from the PD active site (Figs. 3E, F). Although UIS2-NTD lacks structural homologs and shows moderate global confidence (Figs. S8A, B), the α6 helix forms a negatively charged surface (Fig. S8C). Three positively charged arginines (Arg61, Arg62, Arg64) within the S59-loop form a finger-like structure that stabilizes binding by contacting negatively charged residues on UIS2-NTD α6 (Glu164, Asp167, Asp170) (Figs. 3F, G). Additionally, Tyr161 on the UIS2-NTD α6 helix interacts with Ser65 of eIF2α via a hydrogen bond (Figs. S8G–I).

In the second conformation (Fig. 3D), the UIS2-PD active site is exposed, and the S59-loop interacts directly with the catalytic pocket (Fig. 3H). The UIS2-PD alone was predicted with high structural confidence, indicating a well-folded domain (Figs. S8D, E), and features a negatively charged pocket adjacent to the active site (Fig. S8F). In this conformation, the S59-loop interacts electrostatically with UIS2-PD, and Arg64 interacts with the active-site residue Glu774 (Figs. 3I, J; Figs. S8J–L). Together, these models show that UIS2 can recognize P-eIF2α through two distinct electrostatic interfaces, involving either the NTD or the PD, each recognizing the S59-loop.

### UIS2-PD forms a stable substrate-binding pocket with geometry distinct from that of PP1α

AF2-multimer provided insight into the overall binding interface between *Pf* UIS2 and *Pf* eIF2α. To define substrate orientation at the active site, we modeled a complex between *Pf* UIS2-PD and a phosphopeptide from *Pf* eIF2α (residues 55–65, pSer59) using AF3 (Fig. 4A). AF3 placed the pSer59 peptide into a negatively charged pocket near the UIS2-PD active site. We compared this model with the Boltz-1 model.^33^ AF3 yielded an ipTM of 0.65 and a pTM of 0.70 (Figs. S9A–C). Boltz-1 gave a lower ipTM of 0.486 but a similar pTM of 0.713 (Figs. S9D–F), suggesting comparable backbone accuracy but weaker interface confidence. Both methods predicted similar peptide orientation at the active site. The main difference was in residues 618–765, which AF3 predicted as a disordered loop and Boltz-1 as a low-confidence helix. This region lies away from the active site in both models and is unlikely to affect substrate binding or catalysis.

**Fig. 4:**
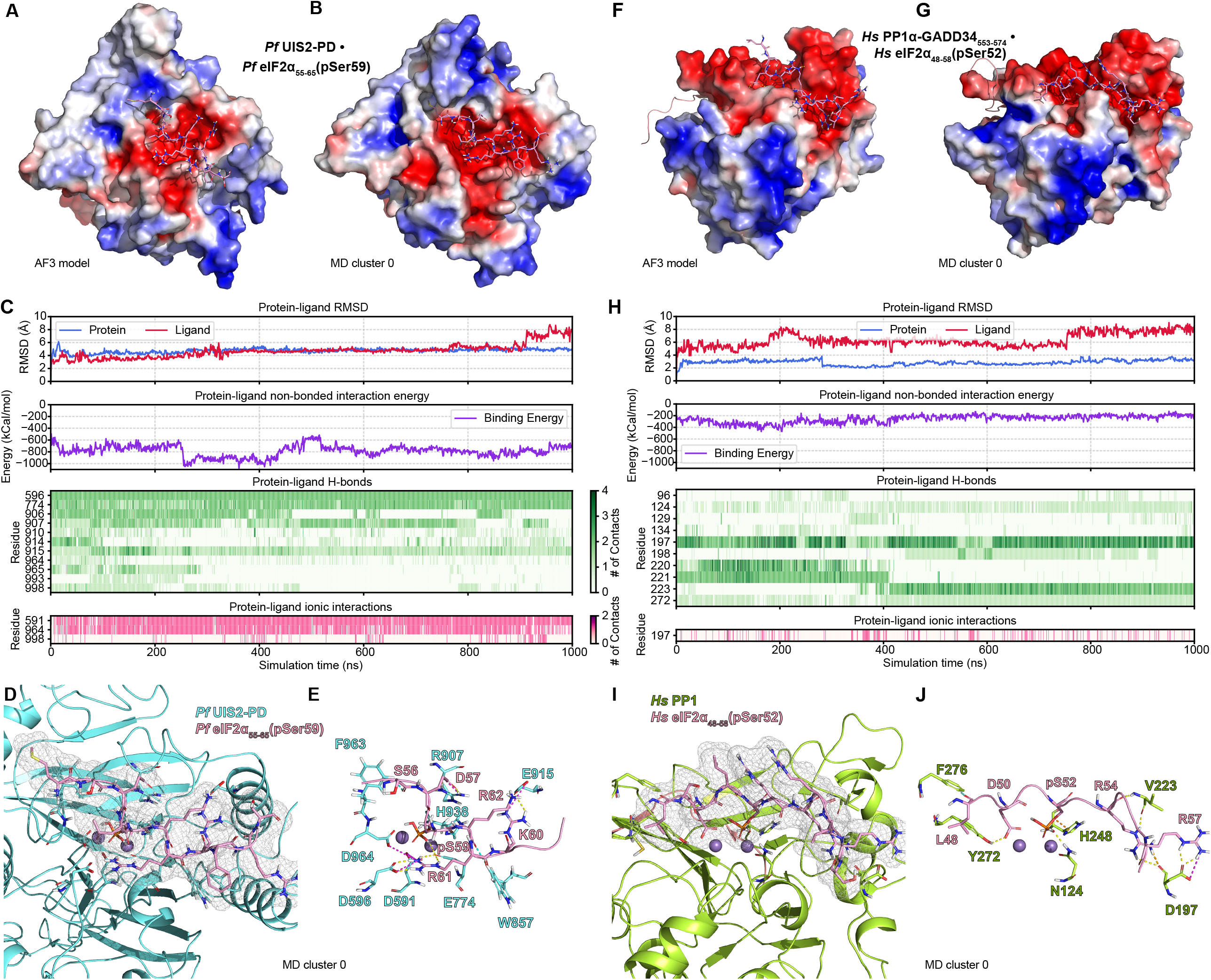
UIS2-PD forms a stable substrate-binding pocket with geometry distinct from that of PP1α. **(A)** Starting structure of *Pf* UIS2-PD bound to the *Pf* eIF2α_55-65_ peptide phosphorylated at Ser59 (pS59), predicted by AlphaFold3 and used to initiate a 1 µs MD simulations. Electrostatic surface of *Pf* UIS2-PD is shown (red, negative; blue, positive), with the *Pf* eIF2α_55-65_(pS59) peptide displayed as a cartoon. **(B)** Cluster 0 representative structure from the MD trajectory of UIS2-PD– eIF2α_55-65_(pS59), corresponding to the most stable binding pose. Trajectory frames were aligned to the UIS2-PD backbone and clustered based on ligand RMSD. Ligand conformations were grouped by orientation within the pocket. **(C)** Time traces from the 1 µs MD simulation: backbone RMSD for UIS2-PD (blue) and ligand RMSD of the pSer59 peptide (red); nonbonded interaction energy (Coulomb + van der Waals) between peptide and UIS2-PD; and heatmaps showing per-residue counts of hydrogen bonds and ionic contacts over time. **(D, E)** Cartoon views of key interactions in the cluster 0 structure. Electron density mesh illustrates geometric complementarity of the pocket around the pSer59 peptide. Color code: yellow, hydrogen bond; magenta, salt bridge; cyan, aromatic hydrogen bond. **(F)** Starting structure of human PP1α in complex with GADD34_553-574_ and the human eIF2α_48-58_ peptide phosphorylated at Ser52 (pS52), predicted by AlphaFold 3 and used for a 1 µs MD simulation. **(G)** Cluster 0 representative structure from the PP1α– eIF2α_48-58_(pSer52) MD trajectory. **(H)** Time traces from the PP1α simulation: backbone RMSD for PP1α (blue) and ligand RMSD for eIF2α(pSer52) (red); nonbonded interaction energy between peptide and PP1α; and heatmaps of per-residue hydrogen bond and ionic contact counts. **(I, J)** Cartoon views of key interactions in the PP1α cluster 0 structure. Electron density mesh highlights peptide fit. Color code as in D–E.

To capture conformational dynamics, we performed a 1 µs molecular dynamics (MD) simulation of the AF3-predicted *Pf* UIS2-PD and eIF2α_55-65_(pSer59) complex in explicit solvent with 0.15 M NaCl. This approach incorporates atomic motion, solvent, and thermal fluctuations to assess binding stability and interactions.^34^ Trajectory frames were aligned to the UIS2-PD backbone and clustered by ligand root mean square deviation (RMSD). Clustering analysis identified three major conformations (cluster 0: Fig. 4B; clusters 1 & 2: Figs. S10A, S10B; overall clustering: Fig. S10C). In all clusters, the phosphopeptide reorients to fit better into the UIS2-PD pocket than in the original AF3 model. The binding pocket remained structurally stable, with only minor side-chain rearrangements observed.

The 1 µs MD trajectory confirmed the complex stability (average peptide RMSD ∼4 Å; Fig. 4C). The eIF2α_55-65_(pSer59) peptide fluctuated for ∼300 ns before settling into a stable binding pose. Nonbonded interaction energies (Coulomb + van der Waals) range from -1000 to -600 kcal/mol, indicating strong affinity. UIS2-PD binding residues form a strong and persistent network of hydrogen bonds and ionic interactions throughout the simulation, particularly at Asp596 and Glu774 (hydrogen bonds), and Asp591 and Asp964 (salt bridges), which likely drive specificity (Fig. 4C). Hydrophobic contacts and water bridges also support peptide positioning, although more transient (Fig. S10C). Cartoon views of representative cluster structures (Figs. 4D–E, S10D–E) illustrate additional residues from the MD trajectory analyses that contribute to substrate anchoring. We found that His938 forms an aromatic hydrogen bond with the pSer59 peptide, stabilizing it near the Mn^2+^ ions for catalysis.

We then compared UIS2 and PP1α binding to their respective eIF2α phosphopeptides. UIS2-PD acts on *Pf* eIF2α-pSer59, while PP1α targets human eIF2α-pSer52. We built AF3 and MD-refined models of PP1α– eIF2α_48-58_(pSer52). Electrostatic maps revealed distinct binding surfaces (Figs. 4F–G). MD trajectory analyses show that PP1α exhibits lower backbone and higher ligand RMSDs than UIS2 (Fig. 4H), indicating a more stable enzyme domain and a more flexible ligand. PP1α–eIF2α_48-58_(pSer52) complexes have higher nonbonded interaction energies (∼ –300 kcal/mol), indicating weaker binding affinity. PP1α forms fewer and less persistent hydrogen bonds and salt bridges than UIS2, although Asp197 forms consistently strong hydrogen bonds with the pSer52 peptide, suggesting a key stabilizing interaction in the PP1α complex. His248 interacts with pSer52 via an aromatic hydrogen bond, positioning the phosphate near the catalytic Mn^2+^ (Figs. 4I–J, S10I-J). This suggests different modes of substrate recognition and stabilization by UIS2 and PP1α.

### UIS2-PD binds its substrate through six cooperative patches, partially targeted by Salubrinal

After identifying the UIS2-PD binding pocket in complex with the eIF2α_55-65_(pSer59) phosphopeptide, we quantified how individual residues contribute to substrate recognition. We ran a 2 µs all-atom MD simulation on the AF3-predicted *Pf* UIS2-PD and eIF2α_55-65_(pSer59) complex under physiological conditions (150 mM NaCl, 300 K, 1 bar). Analysis of the trajectory shows a subset of residues that maintain persistent contacts (> 50 % occupancy) with the peptide (Fig. 5A). These residues cluster into six discrete surface patches (Fig. 5B; Fig. S11A–B). Patch I (D591, D596, E774) lies next to the Mn^2+^ site and anchors Arg61 electrostatically. Patch II (Y777, F778, W857) forms an aromatic, hydrophobic pocket wall, complementing nonpolar or aromatic substrate residues. Patch III (R906, R907) creates a basic ridge that attracts acidic side chains. Patch IV (E914, E915) presents a small acidic surface at the edge that engages basic residues. Patch V (H938) sits centrally and forms a hydrogen bond with pSer59, positioning the phosphate for catalysis. Patch VI (D964, N965, S966, S967) forms a polar surface along the upper groove rim and hydrogen bonds with the peptide. Together, these patches stabilize and orient the peptide for dephosphorylation.

**Fig. 5:**
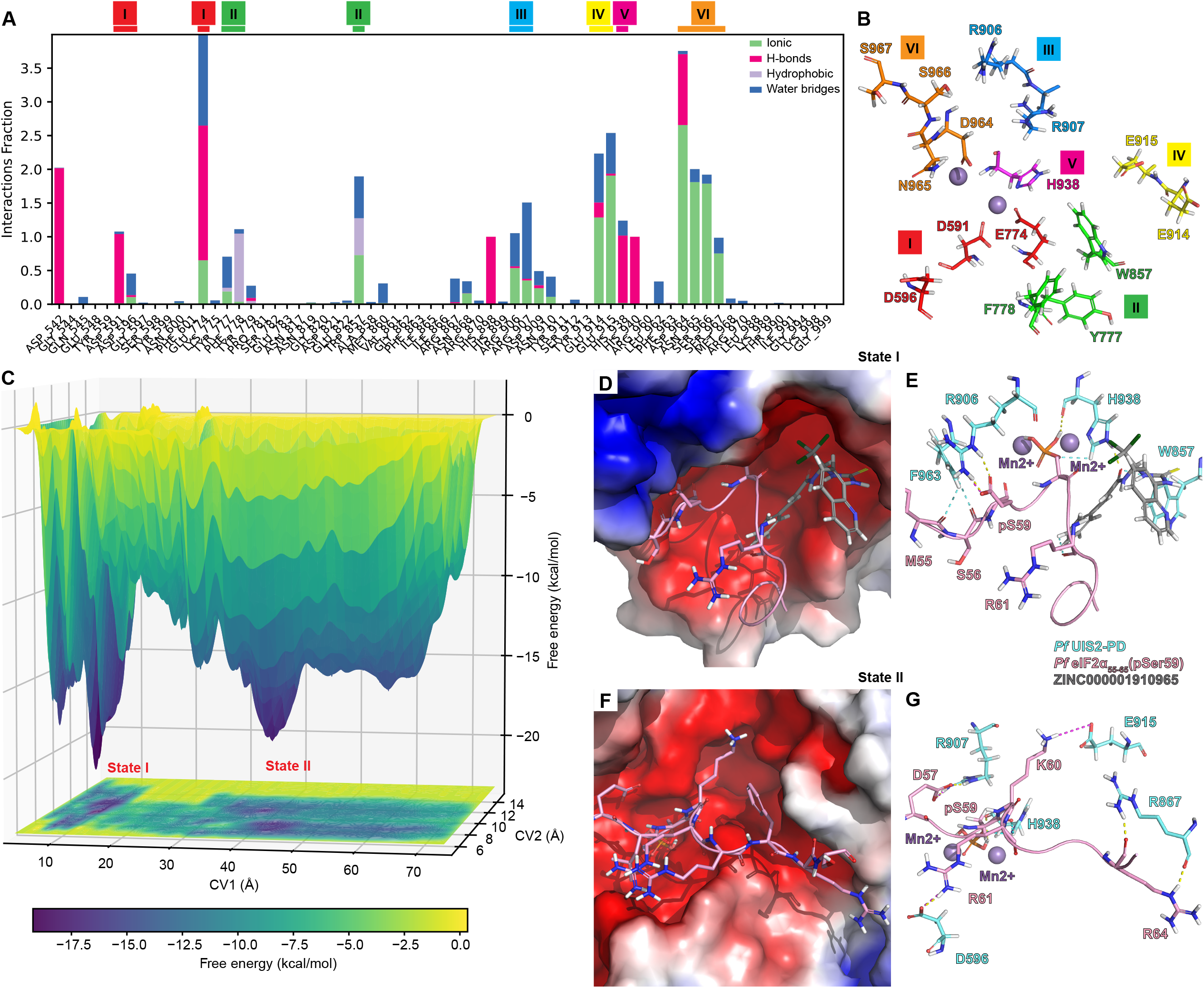
UIS2-PD binds its substrate through six cooperative patches, partially targeted by Salubrinal. **(A)** Persistent peptide-binding residues on UIS2-PD identified from a 2 µs all-atom MD simulation with the eIF2α(pSer59) peptide. Interactions are classified into four categories: hydrogen bonds, hydrophobic contacts, ionic interactions, and water bridges. Stacked bar charts indicate how frequently each interaction occurs throughout the simulation. Values greater than 1.0 reflect multiple same-type contacts formed simultaneously by a single residue. Residues are grouped into six spatially organized patches. **(B)** Spatial clustering of binding patches visualized on the surface of UIS2-PD. Each patch is color-coded and labeled with representative residues. **(C)** 3D metadynamics free-energy surface (200 ns simulation) plotted as a function of CV1 (distance from Salubrinal to UIS2 Cα atoms D591/D596/E774/F778) and CV2 (distance from pSer59 loop backbone to those same residues). Gaussian hills (0.1 kcal/mol height, 0.2 Å width) were added every 1 ps in explicit solvent (300 K, 1.01325 bar, 150 mM NaCl). The yellow-to-purple color scale indicates increasing free energy; valleys represent low-energy binding states. The arrow highlights the two lowest-energy minima. **(D)** Close-up of State I, showing Salubrinal (gray) occupying the sub-pocket normally bound by pSer59. **(E)** Detailed interactions in State I: Salubrinal forms a hydrogen bond with H938 and hydrophobic contact with W857, mirroring Patch II/V engagement of pSer59. **(F)** Alternative low-energy state in which Salubrinal is dissociated and the pSer59 loop is bound to UIS2. **(G–H)** Close-up views of the peptide-only State II, illustrating the interactions that stabilize pSer59 across the binding groove.

To investigate inhibition mechanisms, we performed metadynamics simulations to sample competitive binding between Salubrinal and the pSer59 peptide on UIS2-PD. Standard MD simulations often become trapped in local energy minima and require prohibitively long simulations to sample binding transitions. Metadynamics overcomes this by adding a time-dependent bias to collective variables, lowering energy barriers and accelerating exploration of the binding free-energy landscape. The metadynamics system included *Pf* UIS2-PD, the eIF2α_55-65_(pSer59) peptide, and Salubrinal, solvated under physiological conditions (300 K, 1.01325 bar, 150mM NaCl). We defined two collective variables (CVs): CV1 as the center-of-mass distance from Salubrinal to UIS2 residues D591, D596, E774, F778; CV2 as the distance from the pSer59 loop backbone to the same UIS2 residues. We applied a bias potential and deposited Gaussian hills (0.1 kcal/mol height, 0.2 Å width) every 1 ps to enhance sampling. The accumulated bias was used to reconstruct the 3D free-energy surface (Fig. 5C).

The free-energy plot showed two minima (Fig. 5C), whose depths reflected the thermodynamic stability. The deepest well (State I) corresponds to a ternary complex in which Salubrinal and the eIF2α_55-65_(pSer59) phosphopeptide co-occupy the UIS2 pocket (Figs. S11C, D). Closer inspection (Fig. 5D-E) shows Salubrinal forms a hydrogen bond with H938 and hydrophobic contact with W857, the residues also used in Patch II and V for peptide binding. This overlap confirms that Salubrinal directly competes for the substrate-binding site. State II is the peptide-only complex: the pSer59 peptide remained bound while Salubrinal was dissociated (Figs. S11E, F). The peptide spans the binding groove (Figs. 5G, H), stabilized by the same interactions observed in the MD simulation. State I is ∼3 kcal/mol lower than State II, indicating a thermodynamic advantage for Salubrinal binding. These results demonstrate that Salubrinal competes directly with the pSer59 peptide for UIS2’s active site, inhibiting dephosphorylation.

### Structural and energetic basis of Salubrinal competition at the UIS2 substrate-binding site

To assess Salubrinal’s affinity for *Pf* UIS2-PD and its ability to block substrate rebinding, we used Glide docking and MD simulations. Docking used the MD-relaxed, eIF2α_55-65_(pSer59)-bound conformation of UIS2-PD (Cluster 0 from Fig. 4B). This conformation preserves side-chain orientations essential for ligand recognition and provides a functionally relevant model for competitive binding. Salubrinal contains a thiourea substituted at one nitrogen with a 2-quinolinyl group and at the other with an (S)-2,2,2-trichloro-1-aminoethyl group. The latter group is acylated by the cinnamoyl unit, an α,β-unsaturated amide linked to a benzyl ring (Fig. S12A). Salubrinal binds to the negatively charged UIS2-PD pocket (Fig. 6A). It interacts through three patches: patch III (Arg907), patch IV (Glu915), and patch II (Phe778, Trp857). The thiourea binds Arg907, the secondary amine interacts with Glu915, the benzyl group contacts Phe778, the amide carbonyl forms a π–hydrogen bond with Trp857, and the trichloromethyl group creates a halogen bond with Asn910 (Figs. 6B-C).

**Fig. 6:**
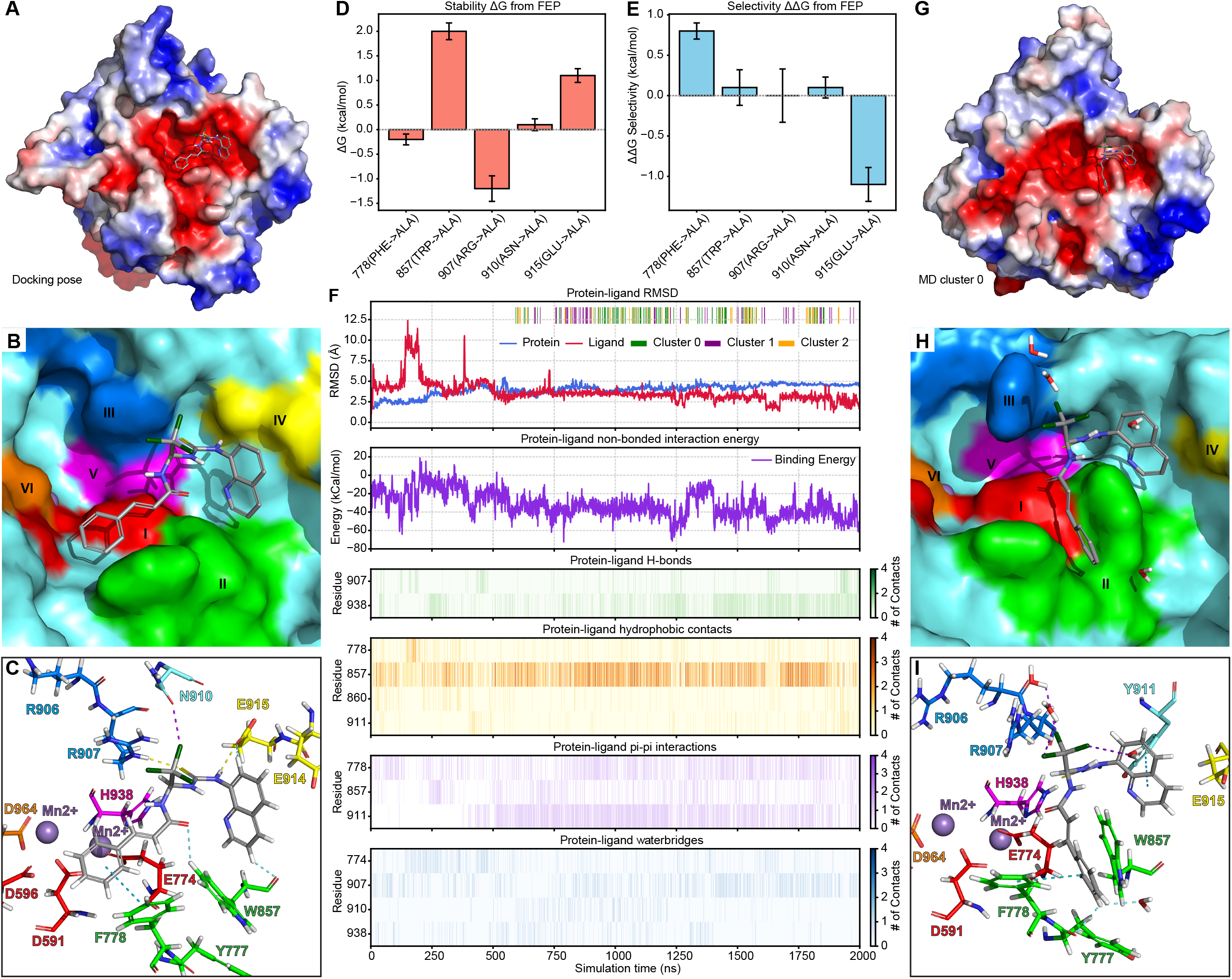
Structural and energetic basis of Salubrinal competition at the UIS2 substrate-binding site. **(A-C)** Glide-predicted docking pose of Salubrinal in the UIS2-PD binding pocket. **(A)** surface view **(B)** Surface representation colored by interaction patches **(C)** Cartoon view showing Salubrinal orientation and key contact residues. **(D–E)** Free energy perturbation (FEP) alanine scanning of five key pocket residues. **(D)** Stability free energy changes (ΔG) quantify individual residue contributions to Salubrinal binding stability. **(E)** Selectivity free energy differences (ΔΔG) for the same residues, quantify their contributions to binding specificity. **(F)** RMSD traces from a 2000 ns MD simulation initiated from the docking pose: blue, UIS2-PD backbone; red, Salubrinal. Cluster assignments are shown below (cluster 0 = green, cluster 1 = purple, cluster 2 = orange). Lower panels show time series of non-bonded interaction energy (Coulomb + van der Waals) and per-residue contact heatmaps, including hydrogen bonds, hydrophobic interactions, π– π stacking, and water-mediated hydrogen bonds. **(G–I)** Representative conformation from MD cluster 0. **(G)** Surface view. **(H)** Surface colored by interaction patches. **(I)** Cartoon view, illustrating the bent ligand conformation in the binding pocket.

We conducted FEP calculations with alanine scanning to quantify residue contributions to Salubrinal binding (Fig. 6D-E). The binding free energy change (ΔG) shows the mutation’s impact on complex stability. Trp857 and Glu915 mutations produce high positive ΔG values (∼1.2–2 kcal/mol), indicating their role in stabilizing the Salubrinal-bound complex. But Phe778, Arg907, and Asn910 mutations show small ΔG changes (∼0 kcal/mol), suggesting a limited role. The relative binding free energy change (ΔΔG) measures the mutation’s effect on ligand binding affinity. Phe778 has a positive ΔΔG (∼0.8 kcal/mol), marking it as crucial for Salubrinal recognition. Yet Trp857 shows near-zero ΔΔG values (∼0 kcal/mol), indicating it contributes more to stability than specificity. Glu915 shows a negative ΔΔG (∼–1.1 kcal/mol), suggesting the mutation improves ligand affinity by relieving strain, despite its stabilizing role. These results show that Trp857 drives complex stability, Phe778 promotes specific binding, and Glu915 balances stabilization with slight inhibition.

To test the stability of the docked pose, we ran 2000 ns of all-atom unbiased MD simulations. The trajectory analyses reveal stable protein-ligand complexes. The ligand conformation stabilizes after ∼500 ns, with RMSD fluctuations maintained between 2-3 Å (Fig. 6F). Non-bonded interaction energy improved from –20 kcal/mol to –40 kcal/mol during the simulation. Hydrogen bonds involving Arg907 and His938 are observed, while hydrophobic contacts and π-π stacking with Phe778, Trp857, and Tyr911 remain strong, supporting their role in ligand binding. Water bridges with Glu774, Arg907, Asn910, and His938 also help strengthen binding. We performed RMSD-based clustering on Salubrinal, identifying three main conformations (Fig. 6F, Cluster 0: Figs. 6G-I; Clusters 1 & 2: Figs. S12B-D, S12E-G). Salubrinal adopts a bent shape in the pocket, with strain around the thiourea and amine linkers. The quinoline ring sits in a hydrophobic subpocket near Tyr911 at patch IV, and polar groups face acidic surface residues. The benzyl group reaches patch II, interacting with Phe778 and Trp857. These findings emphasize cooperative hydrophobic and hydrogen-bonding interactions in Salubrinal binding, offering a structural mechanism for its inhibitory effect on UIS2.

## Discussion

UIS2 is essential for the erythrocytic stage of *Plasmodium*. Its knockout impairs parasite proliferation during this stage (Fig. 1A). We observed strong UIS2 expression in trophozoites (Fig. 1C), a stage marked by active DNA replication and metabolism.^35,36^ During this stage, eIK1 and PK4 phosphorylate eIF2α in response to amino acid deprivation or ER stress, slowing translation and growth.^11,13,17,37^ The co-expression of UIS2, eIF2α, and its kinases (Figs. 1C, E–H) and the reported absence of P-eIF2α in trophozoites,^17^ suggest that UIS2 activity predominates under normal conditions to sustain high protein synthesis. In late schizonts, the parasite completes nuclear division and prepares for egress,^38^ while P-eIF2α remains detectable.^17^ UIS2 transcript levels stay high, and the protein localizes exclusively to the PVM,^8,9^ suggesting that UIS2 is poised to reverse eIF2α phosphorylation and regulate translation at the host–parasite interface. As blood-stage parasites cause clinical symptoms and represent major drug targets, these findings support UIS2 as a therapeutic candidate with potential prophylactic benefits in liver-stage infection.

UIS2-NTD specifically binds to phosphorylated eIF2α, suggesting a regulatory function.^7^ This interaction mechanism differs from that of PP1α, which requires GADD34 to remodel its catalytic site through RVxF and SILK motifs to enable substrate specificity.^20,39-41^ Our AF model illustrates the structural basis of UIS2 interaction (Fig. 2): the pSer59 loop interacts with UIS2-NTD and UIS2-PD through independent electrostatic contacts. The interaction between UIS2-NTD and the pSer59 loop may facilitate spatial recruitment of eIF2α, as the NTD contains a RILDE motif,^8^ separate from α6 that binds P-eIF2α, and likely anchors UIS2 to the PVM. This interaction may also hinder eIF2B binding to P-eIF2α, promoting efficient substrate transfer to UIS2-PD, since the pSer59 loop of eIF2α can also bind the cavity between eIF2Bα and eIF2Bδ subunits of the eIF2B complex, inhibiting its GTP exchange activity.^42-44^ Together, our findings support a spatially coordinated, two-step mechanism in which UIS2-NTD recruits the substrate and UIS2-PD catalyzes its dephosphorylation.

Structural modeling suggests that UIS2-PD confers substrate specificity through its binding pocket, consistent with biochemical evidence that the PD alone is enzymatically active toward *Plasmodium* eIF2α.^7^ The model reveals structural homology between UIS2-PD and PAP5, suggesting comparable substrate recognition strategies. In PAP5, substrate specificity arises from residues that shape the binding pocket to favor substrates such as ATP, PEP, or phytate.^45^ Similarly, MD simulations of UIS2-PD identify six cooperative binding patches that control specificity for the pSer59 loop (Fig. 5A), indicating that UIS2, like PAP5, achieves specificity through the spatial arrangement and chemical properties of residues surrounding the active site. Our results further reveal that substrate sequence alone does not account for binding specificity. Although *Plasmodium* and human eIF2α share sequence similarity in their S1-domain loops, the substrate loops adopt distinct conformations upon binding to their respective phosphatases. These differences reflect variations in binding pocket geometry and interaction networks between UIS2-PD and the PP1α–GADD34 complex. Thus, specificity depends not on the loop sequence, but on side-chain interactions such as hydrogen bonds and ionic contacts that are structurally optimized within each pocket. These structural differences underlie species-specific recognition and offer a basis for designing selective inhibitors.

UIS2-PD exhibits structural features that make it well suited for accurate computational modeling, including a preorganized binding pocket and electrostatic complementarity with the pSer59 loop. The AF-predicted structure of UIS2-PD has a high confidence score (pLDDT > 90), supporting structural accuracy comparable to experimentally determined models.^46^ The UIS2-PD binding pocket remains largely unchanged upon pSer59 loop binding, with only minor side-chain adjustments. MD simulations confirm this structural stability, showing little rearrangement of the binding pocket during phosphopeptide interaction. This binding pocket stability supports a preorganized architecture, similar to that observed in SH2–peptide systems, where AF achieves near-experimental accuracy.^47^ Such preorganization makes UIS2-PD particularly amenable to structure prediction, in contrast to highly dynamic systems like GPCRs, which adopt multiple conformational states,^48^ and reduce reliable modeling.^49,50^

The UIS2-PD–pSer59 loop complex is compatible with MD simulation, allowing conformational refinement beyond the static AF model.^51-53^ AF predicts that UIS2-PD binds the pSer59 loop through electrostatic complementarity and geometric fit. Because electrostatic interactions are sensitive to solvent conditions, MD simulations in explicit solvent serve as a refinement step using physics-based force fields. MD simulations reveal six spatially clustered surface patches on UIS2-PD, formed by residues with similar chemical properties, that ensure persistent pSer59 loop contacts and contribute to substrate specificity. Such dynamic electrostatic interactions are difficult to resolve by NMR due to ensemble averaging, which limits the detection of transient and low-population states.^54^ Together, these features enable high-confidence structural prediction and simulation of UIS2-PD.

Salubrinal acts as a competitive inhibitor for substrate binding in UIS2-PD. Metadynamics and FEP results show that it outcompetes the substrate for PD binding with favorable binding free energy. Salubrinal has been reported to inhibit P-eIF2α dephosphorylation. It also dissociates the GADD34–PP1α complex,^16^ yet drug screening studies dock Salubrinal in the catalytic pocket of PP1α,^55^ not at the GADD34 interface, implying that complex dissociation may result from allosteric effects. MD simulations (Figs. 6G-I, Figs. S12B-G) show that Salubrinal remains stably bound within the UIS2-PD pocket, occupying cooperative binding patches through hydrophobic and hydrogen-bond interactions, and thus blocks substrate rebinding. These findings provide a chemical basis for Salubrinal inhibition. Future inhibitor designs will benefit from experimental validation to expand and refine this structure-based framework.

## Conclusion

Our study shows the critical role of UIS2 in *Plasmodium* development, particularly during the erythrocytic stages. The structural and mechanistic basis of UIS2-mediated eIF2α dephosphorylation involves a two-step substrate recognition mechanism. UIS2-PD contains a conserved, stable architecture that binds the S59-containing loop of *Plasmodium* eIF2α through electrostatic interactions distinct from those of human PP1α. Salubrinal competitively inhibits this binding through hydrophobic and hydrogen-bond interactions. Our work establishes UIS2 as a biologically essential enzyme and a structurally defined target for antimalarial drug development.

## Methods

### Essential gene analysis and single-cell gene expression analysis

The phenotype screening data were sourced from PlasmoGEM.^21^ To create the phenotype plot, the scatterplot() function from the Python package “seaborn” was used, with translation initiation-related genes annotated from the phenotype dataset. Single-cell transcriptome profiles of 1,787 cells covering all life cycle stages were obtained from the Malaria Cell Atlas (dataset name: *P. berghei* SS2 set1). Stage annotation data were extracted from “pb-ss2-set1-ss2-data.csv,” and single-cell RNA-seq data from “pb-ss2-set1-ss2-exp.csv.” UMAP coordinates for each cell, provided in the original publication,^22^ were used as given in the “pb-ss2-set1-ss2-data.csv” file. Cells were colored based on gene expression values from the “pb-ss2-set1-ss2-exp.csv” file. Data selection and merging were performed using the Python package “pandas,” and plots were generated using the scatterplot() function from the “seaborn” package.

### Structural template search and multiple sequence alignment

HHsearch (v3.3.0) from AF2.3 was used to perform a structural template search against the PDB70 database. This search generated the pdb_hits.hhr file, which contains profile–profile alignments between PfUIS2 and structural templates in the Protein Data Bank. HHsearch compares hidden Markov models (HMMs) to identify remote homologs, ranking results based on probability scores and E-values.

Multiple sequence alignments (MSAs) were generated using Clustal Omega with default settings. The alignment included the following protein sequences: *Pb* UIS2 (A0A509AQ20), *Pf* UIS2 (Q8IKJ1), *Hs* PAP5 (P13686), *Pf* GAP50 (Q8I2×3), *Pb* eIF2alpha (A0A509AJP9), *Pf* eIF2alpha (Q8IBH7) and *Hs* eIF2α (P05198). MSA results were visualized using ESPript 3.0, which was configured to display predicted secondary structure features derived from the AF *Pf* UIS2 model and to highlight conserved sequence motifs. Sequence logos were generated to visualize residue conservation across orthologs, with similarity scores calculated using the Risler substitution matrix.

### Protein structure modeling

For the full-length Pf UIS2 structure prediction, both AlphaFold 2.3 (AF2.3, monomer mode) and AF3 were evaluated and yielded highly similar results. For large protein–protein complexes, including full-length *Pf* UIS2 bound to *Pf* eIF2α, we used AlphaFold-Multimer v2.3 (AF-M2.3).^32^ AF-M2.3 uses deep multiple sequence alignments (MSAs) to capture co-evolutionary signals, which yields more accurate models for multi-domain assemblies.^56,57^ Comparing AF-M2.3 and AF3 models, we found that AF-M2.3 more accurately recapitulated published biochemical data, especially at the interface between the UIS2 N-terminal domain and the eIF2α S1 loop. For receptor–peptide complexes, including UIS2–eIF2α_55-65_(pSer59) and PP1α–eIF2α_48-58_(pSer52), we used AF3 that supports explicit modeling of phosphorylated serine and bound Mn^2+^. AF3 uses simplified MSA processing with a diffusion-based structure module, which improves accuracy for short peptide interactions.^46^ All AF-M2.3 calculations were run on the Harvard Medical School HPC cluster, using either NVIDIA A100 (80 GB VRAM) or RTX 8000 (48 GB VRAM) GPUs with 80 GB RAM reserved per job. We set db_preset=full_dbs and max_template_date=2022-01-01. AF3 predictions were performed on the official AlphaFold server, and Boltz-1 models were generated via the Neurosnap webserver.^33^ Amino acid sequences were obtained from the UniProt database: *Pf* UIS2 (Q8IKJ1), *Pb* UIS2 (A0A509AQ20), *Pf* eIF2α (Q8IBH7), *Pb* eIF2α (A0A509AJP9), *Hs* PP1α (O75807), *Hs* eIF2α (P05198) and *Hs* GADD34 (O75807). All AlphaFold predictions were performed with default parameters, and the highest-confidence models (ranked by predicted LDDT and PAE) were retained for further analysis. The resulting AlphaFold models were subsequently relaxed and evaluated for consistency with available experimental data. The highest-confidence models, ranked by predicted LDDT and PAE, were Amber-relaxed. The top-ranked model for each task was selected for illustration using PyMOL3. All non-Alphafold calculations were conducted on a System76 “Serval” mobile workstation with an Intel Core i7-8700k, running Pop!_OS 22.04 LTS.^58^

### Protein structure preparation

The protein structures of UIS2–eIF2α_55-65_(pSer59) and PP1α–eIF2α_48-58_(pSer52) were prepared using Schrödinger’s PrepWizard (Schrödinger Release 2025-1). The input structures were processed with the following settings: side chains were filled (−fillsidechains), disulfide bonds were detected (−disulfides), termini were capped with neutral acetyl and N-methylamide groups (−captermini), and all protonation states and tautomers were assigned (−assign_all_residues, -include_epik_states). Histidine protonation and residue re-treatment were handled automatically (−rehtreat). Epik was run at pH 7.4 with a tolerance of ±2.0 pH units (−epik_pH 7.4 -epik_pHt 2.0), and PROPKA was used to refine pKa estimates at pH 7.4 (−propka_pH 7.4). A single state per residue was retained (−max_states 1). Water sampling was enabled within 5 Å of the peptide surface (−samplewater -watdist 5.0), and Epik-generated states were included in the output ensemble. The OPLS_2005 force field was applied for all structural refinements (−f OPLS_2005), and conformers were clustered with a 0.3 Å RMSD cutoff (−rmsd 0.3).

### System setup and molecular dynamics simulations

We used Schrödinger’s multisim workflow to build and solvate the protein structures. Protonation states were assigned at pH 7.4 using PROPKA. The System Builder stage employed an orthorhombic box with a 10 Å buffer distance between the solute structures and the simulation boundary box. SPC water was added to fill the orthorhombic box. The system was neutralized with Cl− counterions and supplemented with 0.15 M NaCl (Na+/Cl−) to mimic physiological ionic strength. The OPLS_2005 force field was applied at both the build_geometry and assign_forcefield stages.

All MD simulations were performed with Schrödinger Desmond v2025-1 using an orthorhombic box defined in the system setup and periodic boundary conditions. Protonation states were assigned at pH 7.4 via Epik/PROPKA, and OPLS_2005 parameters were applied to protein, peptide, water, and ions. The system was solvated with SPC water, neutralized with Cl−, and adjusted to 0.15 M NaCl. Equilibration comprised five sequential MD stages, each with restraints on solute heavy atoms unless stated otherwise. We began with Brownian dynamics NVT at 10 K for 100 ps (timestep 1 fs; heavy-atom restraints 50 kcal mol^−1^ Å^−2^). This was followed by NVT at 10 K for 12 ps (Langevin thermostat, τ = 0.1 ps), then two NPT steps at 10 K for 12 ps each (Langevin piston barostat, τ = 50 ps), the first retaining and the second releasing heavy-atom restraints. Finally, we heated the system to 300 K in NPT for 24 ps (MTK barostat, τ = 2 ps; Nose– Hoover thermostat, τ = 1 ps) with no restraints. Nonbonded cutoffs were 9 Å throughout, and the RESPA integrator used 2 fs inner and 6 fs outer timesteps. Production runs in NPT (300 K, 1.013 bar) employed the Multigrator integrator with the same timestep scheme and cutoffs. Each trajectory lasted 1 µs or 1 µs, with frames saved every 1 ps.

### Simulation trajectory analysis

Trajectories were processed using Schrödinger’s Simulation Interaction Diagram to extract RMSD, binding energy, and protein–ligand contact data. Conformational clustering was performed with Schrödinger’s trj_cluster.py, sampling frames every 1 ps and specifying three clusters (−n 3). Clustering was based on ligand heavy-atom RMSD after alignment of each frame to a reference structure using the protein backbone atoms. The MD output CMS file and its trajectory directory served as inputs. This step generated separate CMS files and trajectory subdirectories for each cluster. Per-cluster coordinates and frame assignments were then extracted with trj2mae.py, producing MAE snapshots and CSV tables of frame indices. A second clustering pass (without -split-trj) yielded one representative frame per cluster. All commands were executed via $SCHRODINGER/run in an SBGrid environment. Representative structures were inspected in Maestro for downstream analyses.

Data integration and visualization used a custom Python pipeline. Pandas loaded RMSD, energy, contact, and cluster CSV files. Heatmap matrices for hydrogen bonds, hydrophobic contacts, halogen bonds, water bridges, and π–π interactions were computed by grouping frame–residue counts and pivoting into matrices. Cluster spans were overlaid on the RMSD plot as colored bars.

### Metadynamics

Metadynamics was performed in Desmond v2024-4. We applied the OPLS_2005 force field and solvated the system with SPC water in an orthorhombic box with a 10 Å buffer. Sodium and chloride ions were added to 0.15 M and to neutralize the system. The simulation system included *Pf* UIS2-PD, Salubrinal, and the pS59 peptide. Two distance-based collective variables (CVs) were defined between the ligand heavy-atom group and UIS2 residues 591–596 and 774–778. Gaussian hills of height 0.1 kcal mol^−1^ and width 0.2 Å were deposited every 1 ps over a 200 ns production run. A bias factor tempered the added potential.

All simulations used the NPT ensemble with the MTK barostat (τ = 2 ps) and Nose–Hoover thermostat (τ = 1 ps) at 300 K and 1 bar. The Multigrator integrator employed a 2 fs inner timestep and 6 fs outer timestep under RESPA. A 9 Å cutoff was applied to van der Waals and short-range electrostatics, and long-range interactions were treated via the Useries method. The CV sequence was recorded at the same interval as the trajectory. Free-energy surfaces along the two CVs were reconstructed using Desmond’s metadynamics analysis tools. Representative free energy minima and transition conformations were visualized to interpret ligand binding mechanisms.

### Ligand preparation and Glide docking

Salubrinal structure was downloaded from the Zinc database. LigPrep (Schrödinger v2025-1) was used to prepare Salubrinal. Hydrogens were added and tautomers were generated with Epik at pH 7.4 ± 0.0, followed by desalting, neutralization, and chiral specification. Minimization used the OPLS_2005 force field.

Docking grids were prepared using Glide v2025-1 (mmshare b39c0a02be). Protein atom types and parameters were assigned with OPLS_2005. Van der Waals radii for receptor atoms were scaled by 1.0, and a polarity charge cutoff of 0.25 was applied. Grid dimensions were set to encompass the UIS2-PD and pSer59 peptide interface (nsites = 125) and a buffer size of 1.0 Å. Docking results were further subjected to MD simulations.

### FEP with alanine scanning

Free energy perturbation (FEP) calculations were performed using the Schrödinger FEP Web Service (Schrödinger Release 2025-1). Single-residue alanine scanning mutations were modeled using a dual-topology approach. All systems were prepared using the default protocol provided by Schrödinger. The OPLS5 force field was applied to all simulations. FEP simulations were conducted using 12 λ-windows with 5.0 ns of molecular dynamics sampling per window. Replica exchange with solute tempering (REST2) was employed to enhance sampling around the mutated residues. Stability free energy changes (ΔG) and selectivity free energy differences (ΔΔG) were calculated using the Bennett acceptance ratio (BAR) method. Convergence was assessed based on cycle closure errors and energy convergence quality metrics provided by the FEP Web Service.

## Data availability

The python scripts used for data analysis and figure generation are available at: (https://github.com/a3609640/UIS2).

The PDB data for protein structure modeling and simulation trajectories have been submitted to Zenodo.

## Author Contributions

Su Wu: Conceptualization, Data curation, Formal analysis, Software, Visualization, Writing. Gerhard Wagner: Conceptualization, Funding acquisition, Supervision, Review.

## Acknowledgments

This work was supported by the National Institute of Allergy and Infectious Diseases (5P01AI143565-03, to G.W.). We thank Drs. Philipp Aschauer, Christoph Gorgulla, and Sorin Draga for their valuable scientific discussions.

