## Supplemental information for "Mechanistic Insights into the *Plasmodium* Phosphatase UIS2 in eIF2α Dephosphorylation via Integrated Structure-Based Modeling"

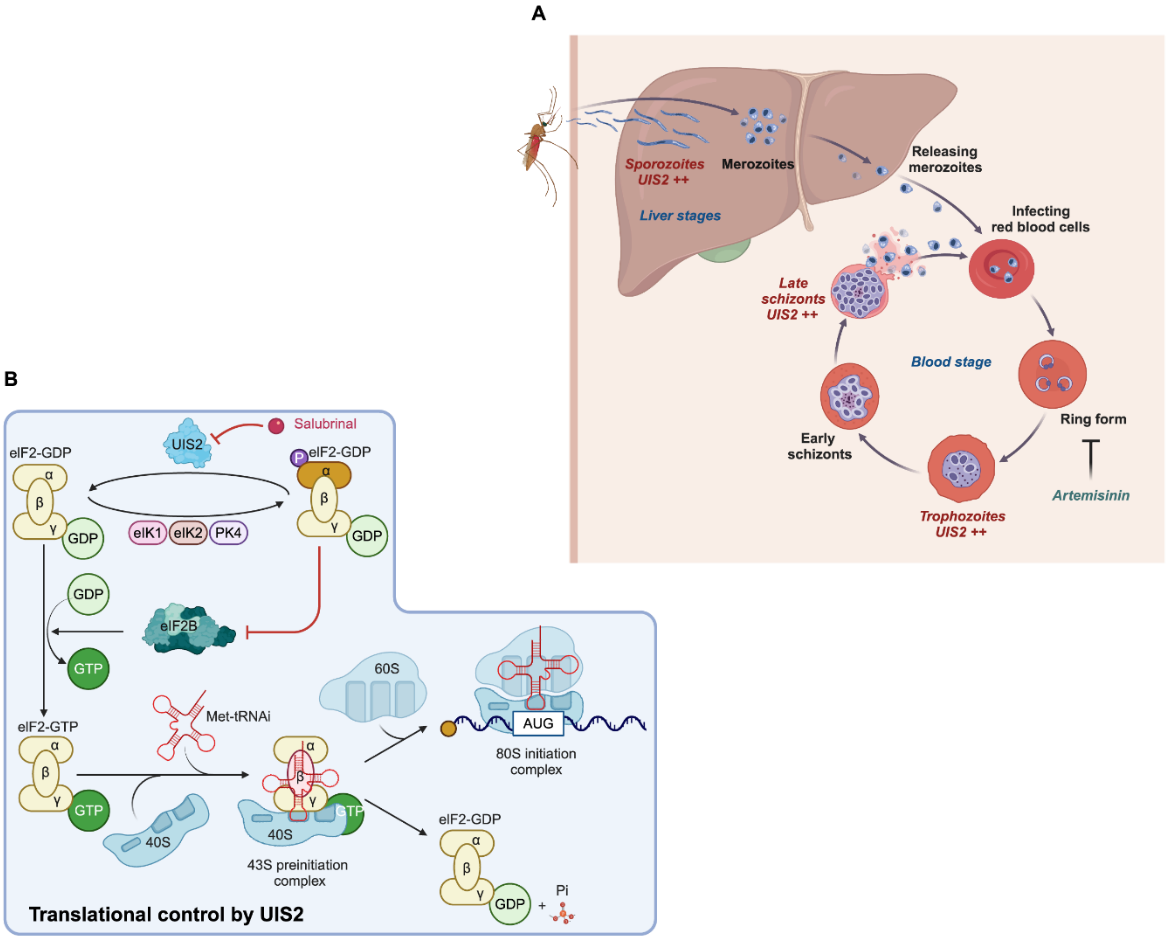


**Supplementary Figure 1. *Plasmodium* life cycle and UIS2 function in translation initiation. (A)** The diagram shows the *Plasmodium* life cycle in the human host, including hepatic and erythrocytic stages. Stages with elevated UIS2 expression are marked in red. **(B)** The schematic illustrates UIS2's role in dephosphorylating eIF2α to promote translation initiation.

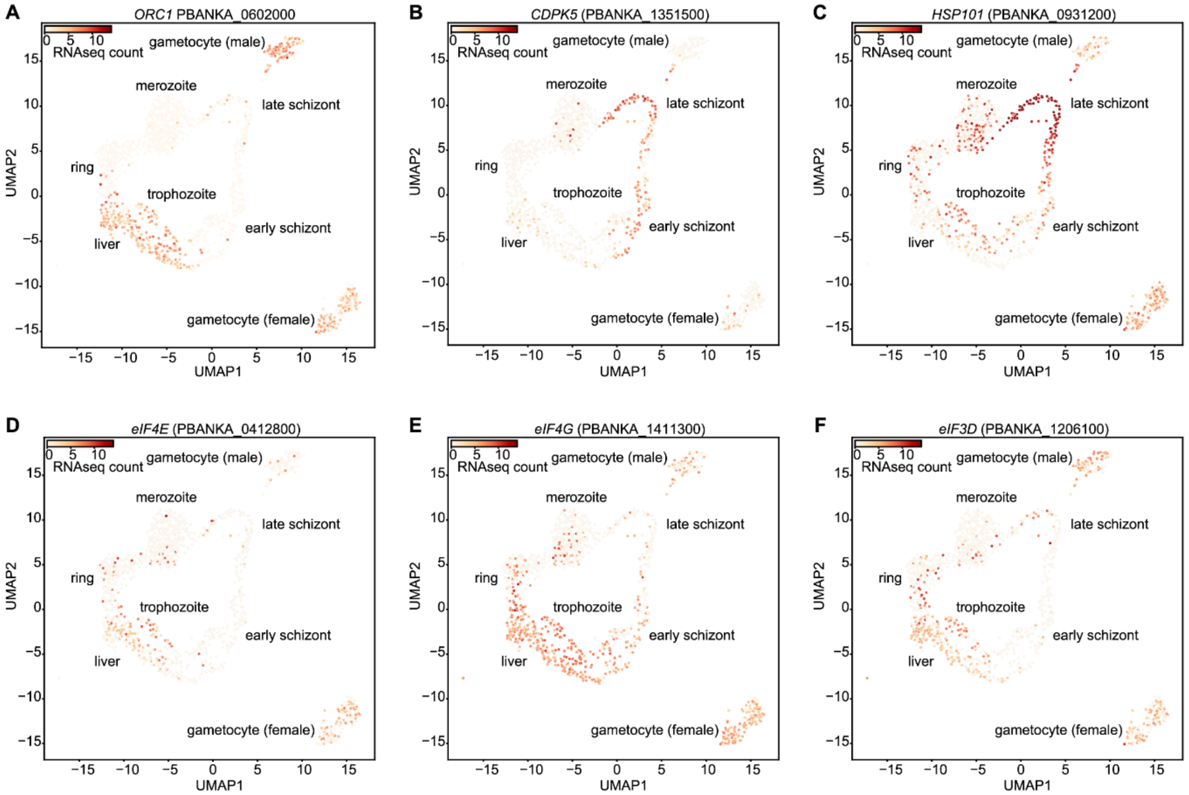


**Supplementary Figure 2. Expression of eIF4F complex components in the liver and trophozoite stages of parasites within the mouse host. (A–C)** UMAP plots show the expression of marker genes for the liver, trophozoite, and late schizont stages in single-cell RNA-seq data from *Plasmodium berghei*-infected mice. **(D–F)** UMAP plots display the expression patterns of genes encoding eIF4F complex subunits (eIF4E, eIF4G, and eIF4A), which are enriched in the liver and trophozoite stages.

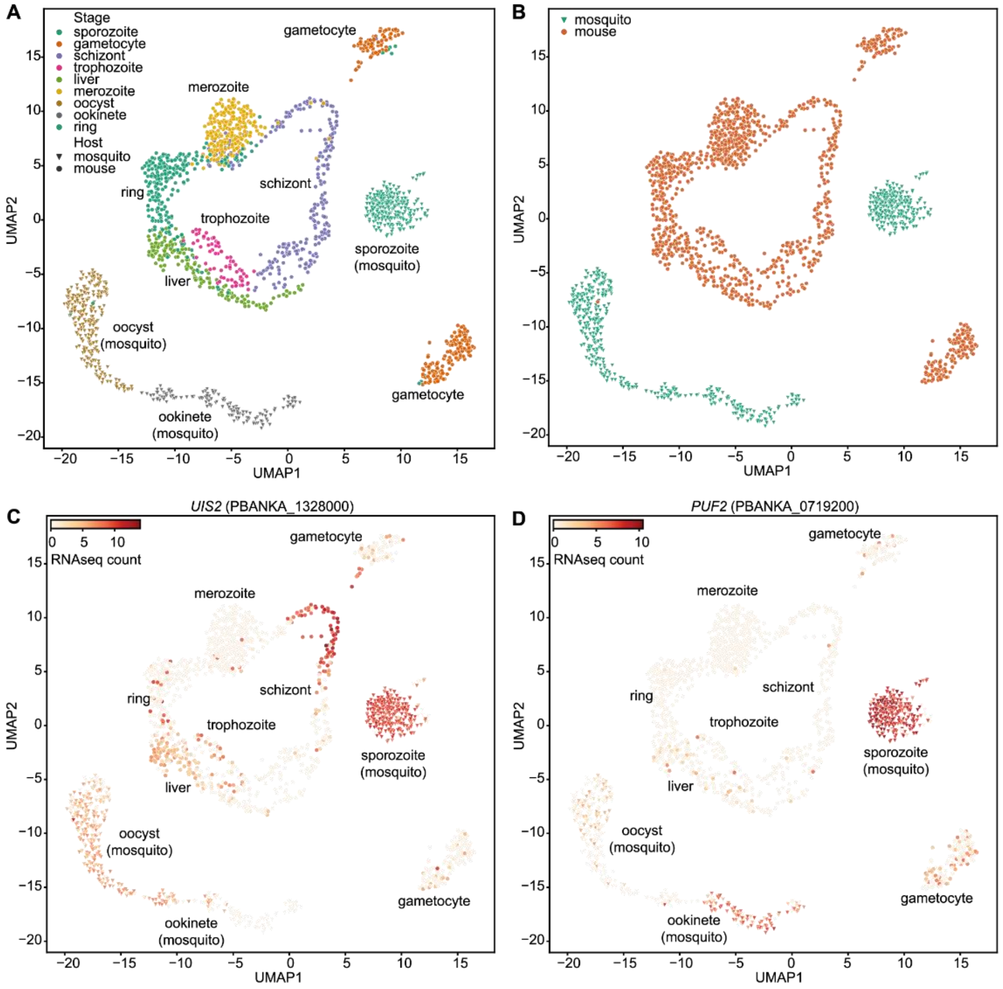


**Supplementary Figure 3. UIS2 translation repression in mosquito-hosted parasite stages. (A)** The UMAP plot shows 1,787 *P. berghei* parasites across all life cycle stages, clustered by transcriptome similarity. Parasites from each stage were isolated and pooled for single-cell RNA-seq. Each dot represents a single parasite, colored by developmental stage. **(B)** The UMAP plot from **(A)** is colored by the host (mouse or mosquito) from which parasites were isolated. **(C–D)** UMAP plots show expression of UIS2 and Puf2 across parasite life stages. Parasites expressing each gene are in red, with shading intensity corresponding to relative transcript abundance in the single-cell RNA-seq dataset.

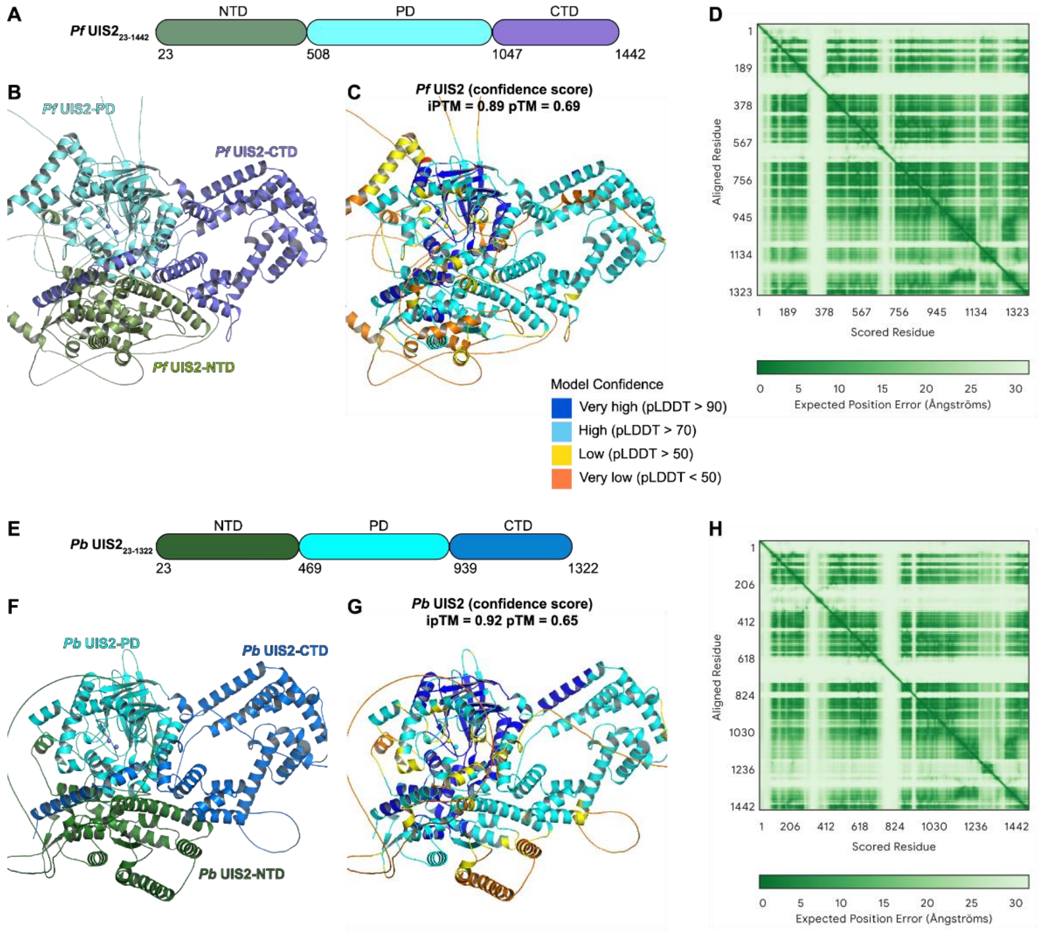


**Supplementary Figure 4. Alphafold predicts similar structures for UIS2 from *P. berghei* and *P. falciparum* with high confidence in the phosphatase domains. (A)** Schematic representation of *Pf* UIS2 domains, including the signal peptide (SP), N-terminal domain (NTD), phosphatase domain (PD), and C-terminal domain (CTD). **(B)** Full-length structural model of *Pf* UIS2 (residues 1–1442) predicted by AlphaFold3. **(C)** Residue-level confidence scores (pLDDT) are mapped onto the *Pf* UIS2 model, with color coding indicating model certainty. **(D)** Predicted aligned error (PAE) plot for *Pf* UIS2, indicating the expected positional error (in Å) between residue pairs. Lower error values (green) correspond to higher structural confidence. **(E)** Schematic diagram of *Pb* UIS2 domains. **(F)** AlphaFold3-predicted structure of full-length *Pb* UIS2 (residues 1-1322). **(G)** Residue-level confidence (pLDDT) scores are color-coded on the model. **(H)** PAE plot for *Pb* UIS2, illustrating residue-wise alignment error across the full-length structure.

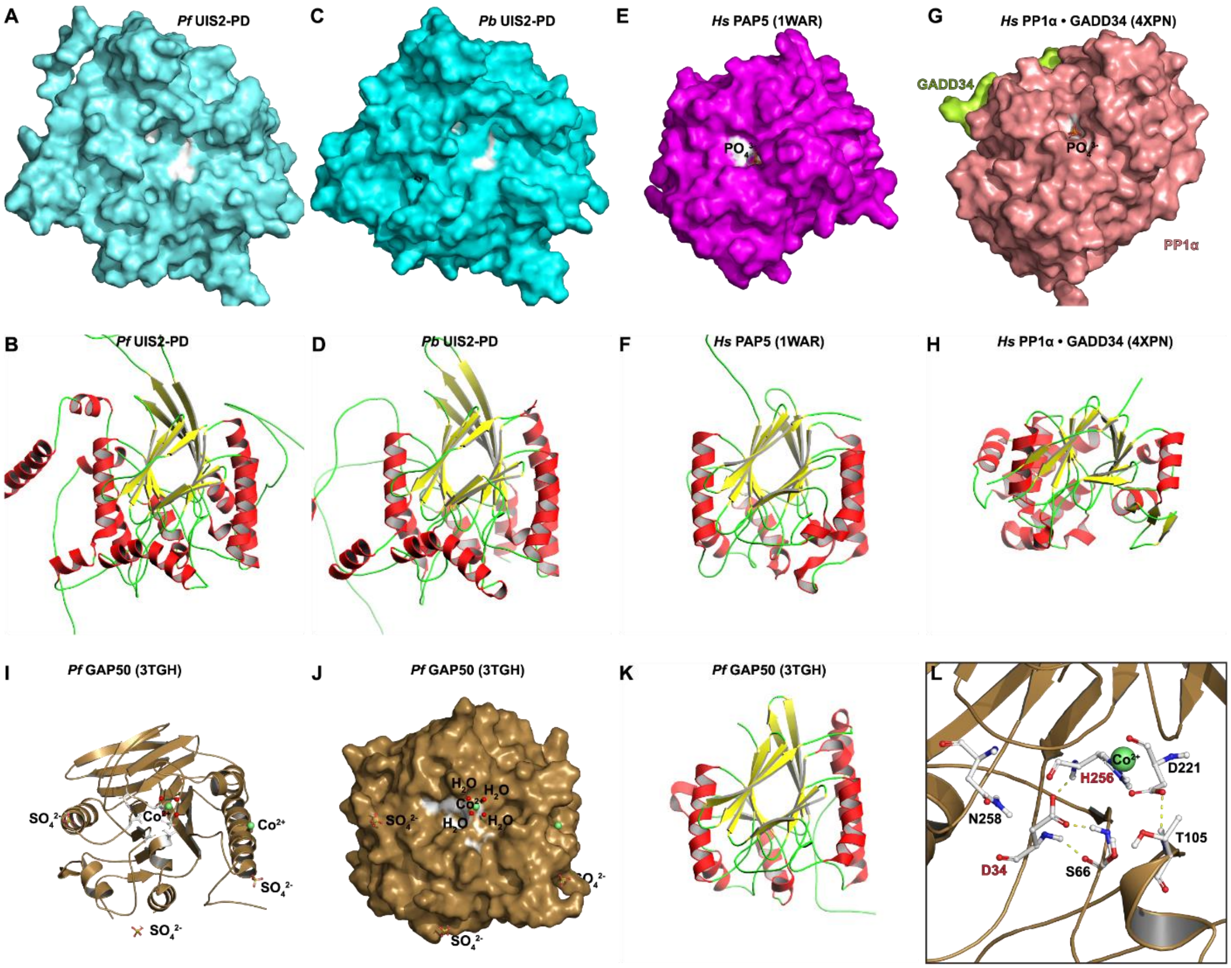


**Supplementary Figure 5. The phosphatase domain of UIS2 shares greater structural similarity with PAP5 than with PP1α. (A-B)** Surface and ribbon representations of *Pf* UIS2-PD. Six conserved active-site residues are shown in white to mark the catalytic cleft. Secondary structural elements are color-coded: a-helices in red and b-strands in yellow. **(C-D)** Surface and ribbon representations of *Pb* UIS2-PD. **(E-F)** Surface and ribbon representations of *Hs* PAP5 (PDB: 1WAR). **(G-H)** Surface and ribbon representations of human PP1α in complex with its regulatory subunit GADD34 (PDB: 4XPN). **(I-J)** Cartoon and surface representation of *Pf* GAP50. The cobalt ions are shown as green spheres and water molecules as red spheres. **(K-L)** Ribbon representation and close-up view of *Pf* GAP50, with the residues labeled at the putative active site.

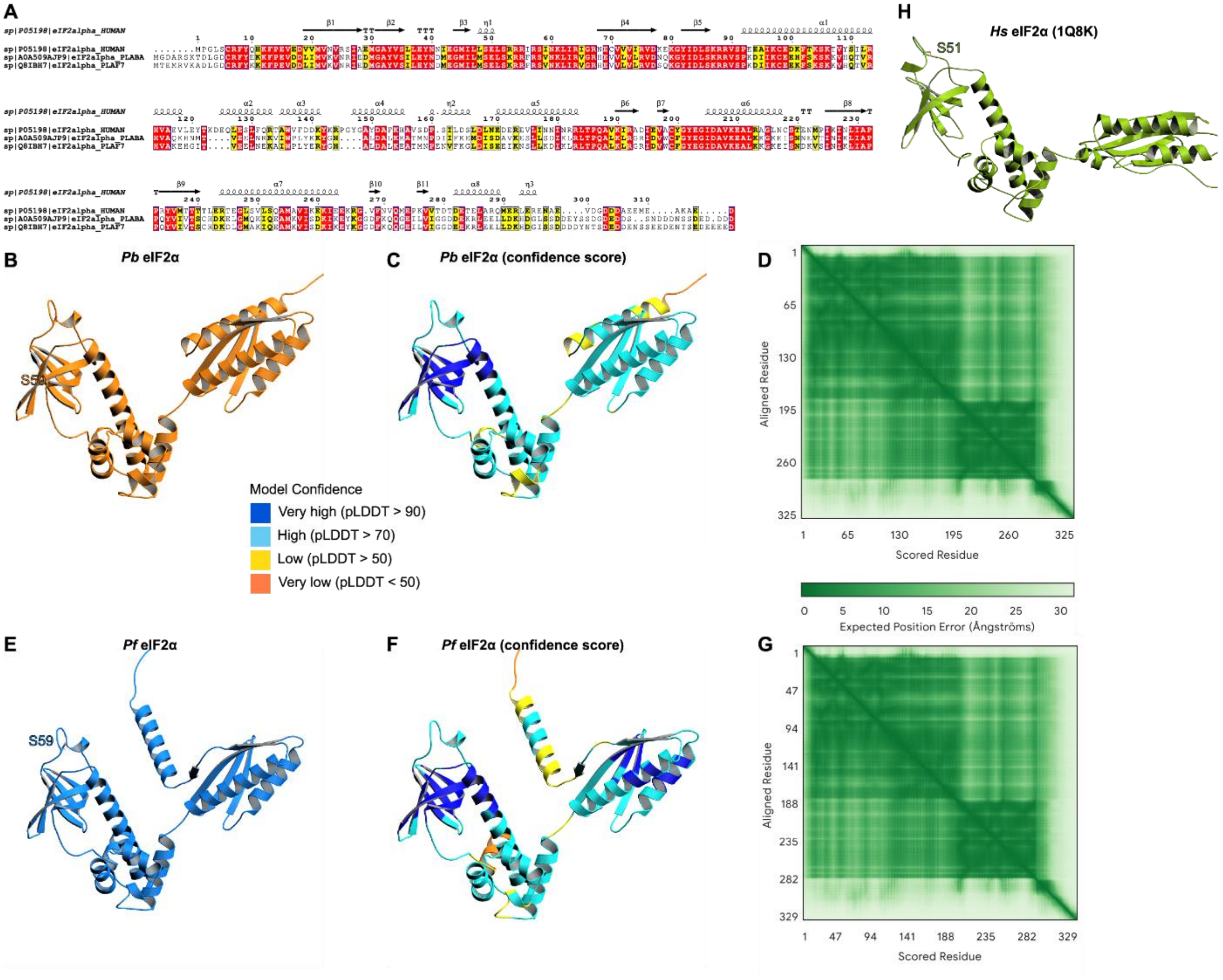


**Supplementary Figure 6. Sequence and structural homology of human, *P. berghei* (*Pb*), and *P. falciparum* (*Pf*) eIF2α. (A)** Multiple sequence alignment of human, *Pb*, and *Pf* eIF2α using Clustal Omega and ESPript 3.0. Secondary structure elements: a-helices (squiggles), $3_{10}$-helices (small squiggles), b-strands (arrows), b-turns (TT letters). Conserved residues are in red. **(B)** Cartoon model of *Pb* eIF2α predicted by AlphaFold 2.3 monomer. **(C)** *Pb* eIF2α model color-coded by pLDDT score to indicate per-residue confidence. **(D)** Predicted Aligned Error (PAE) plot for *Pb* eIF2α, showing the expected positional error (in Å) between residue pairs. **(E)** Cartoon model of *Pf* eIF2α predicted by AlphaFold 2.3 monomer. **(F)** *Pf* eIF2α model colored by pLDDT confidence. **(G)** Predicted Aligned Error (PAE) plot for *Pf* eIF2α. **(H)** Solution structure of human eIF2α determined by NMR (PDB: 1Q8K).

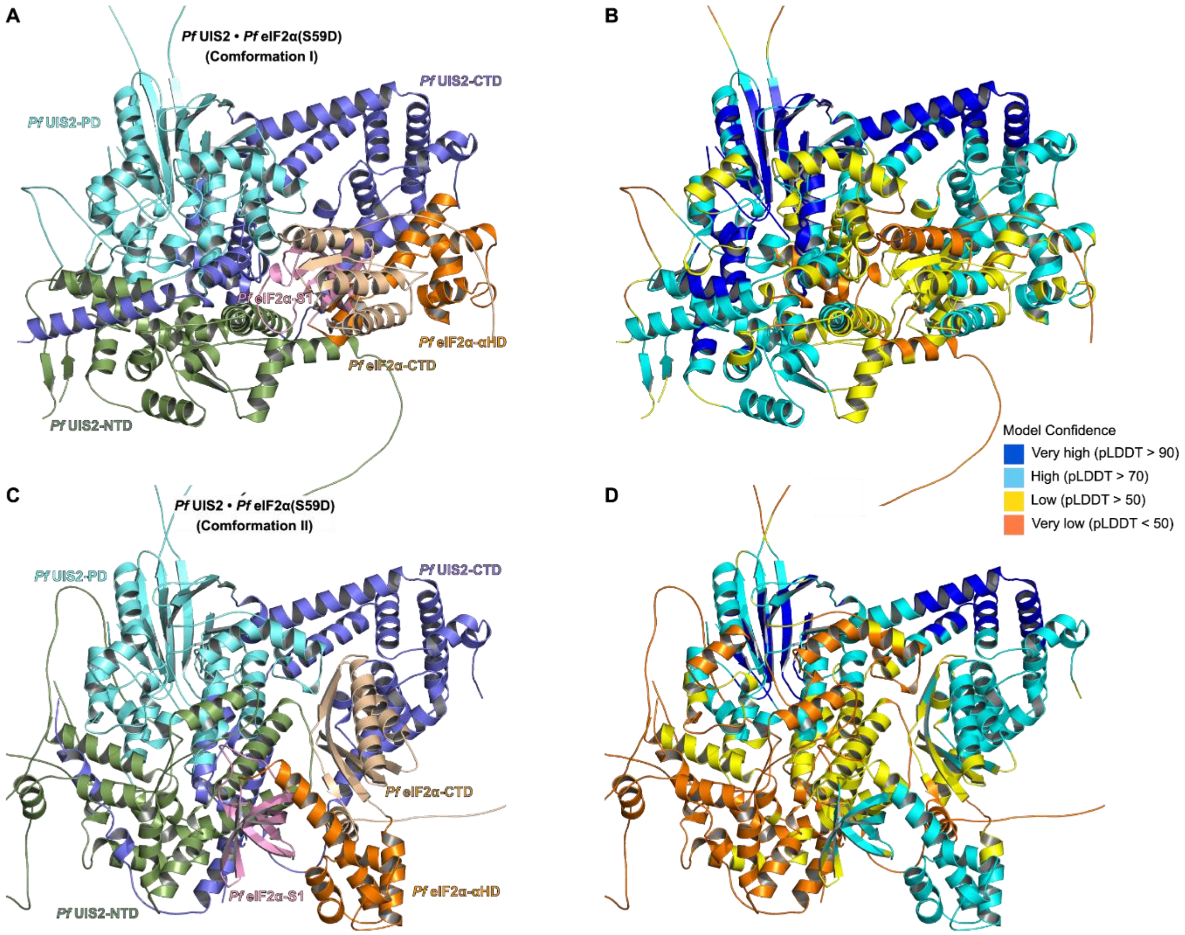


**Supplementary Figure 7. Predicted complex structures of *Pf* UIS2 bound to *Pf* eIF2αS59D**. **(A)** Predicted complex structure of *Pf* UIS2 bound to *Pf* eIF2αS59D in Conformation I, generated using AlphaFold v2.3 multimer. **(B)** Conformation I model colored by pLDDT scores to indicate per-residue confidence. **(C)** Predicted complex structure of PfUIS2 bound to Pf eIF2α S59D in Conformation II, also generated using AlphaFold v2.3 multimer. **(D)** Conformation II model colored by pLDDT confidence.

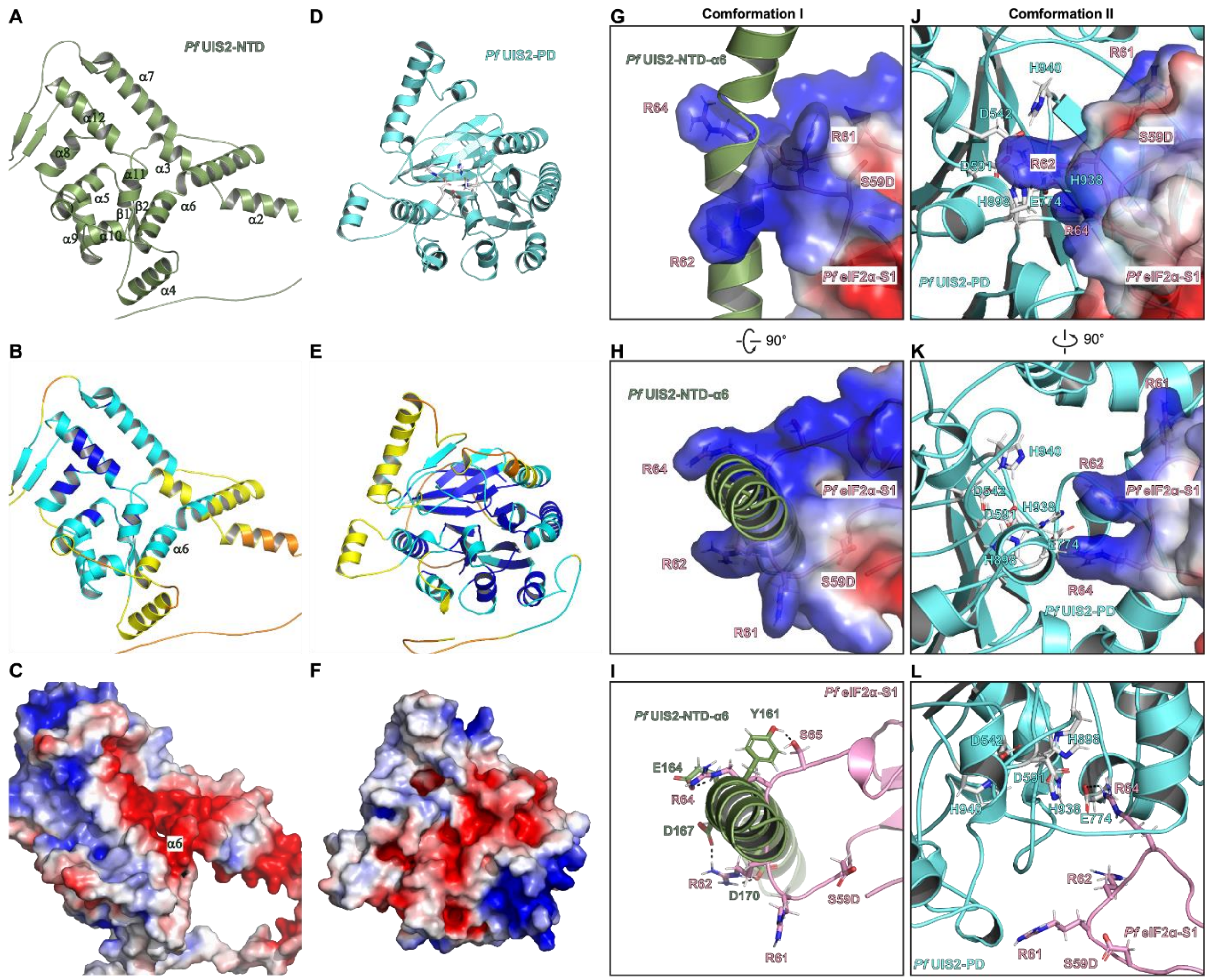


**Supplementary Figure 8. Interaction of the S59-loop of *Pf* eIF2α S59D with helix α6 from *Pf* UIS2-NTD or *Pf* UIS2-PD in two conformations. (A-C)** *Pf* UIS2-NTD monomer model predicted by AlphaFold v2.3. **(A)** Cartoon representation of *Pf* UIS2-NTD. **(B)** *Pf* UIS2-NTD colored by pLDDT confidence score. **(C)** Electrostatic surface of *Pf* UIS2-NTD (red: negative charges; blue: positive charges). **(D-F)** AlphaFold v2.3 monomer model of *Pf* UIS2-PD, with key catalytic residues shown as silver sticks. **(D)** Cartoon model of *Pf* UIS2-PD. **(E)** *Pf* UIS2-PD colored by pLDDT confidence score. **(F)** Electrostatic surface of *Pf* UIS2-PD (red: negative charges; blue: positive charges). **(G-I)** Close-up views of *Pf* UIS2 and *Pf* eIF2α S59D complex (conformation I) predicted by AlphaFold 2.3 multimer, showing helix α6 of *Pf* UIS2-NTD (electrostatic surface; red: negative; blue: positive) interacting with the S59-loop of *Pf* eIF2α S59D (cartoon). Polar contacts are indicated by dashed lines. **(J-K)** Close-up views of *Pf* UIS2 and *Pf* eIF2α S59D complex (conformation II), highlighting *Pf* UIS2 PD (electrostatic surface; red: negative; blue: positive) interaction with the S59-loop.

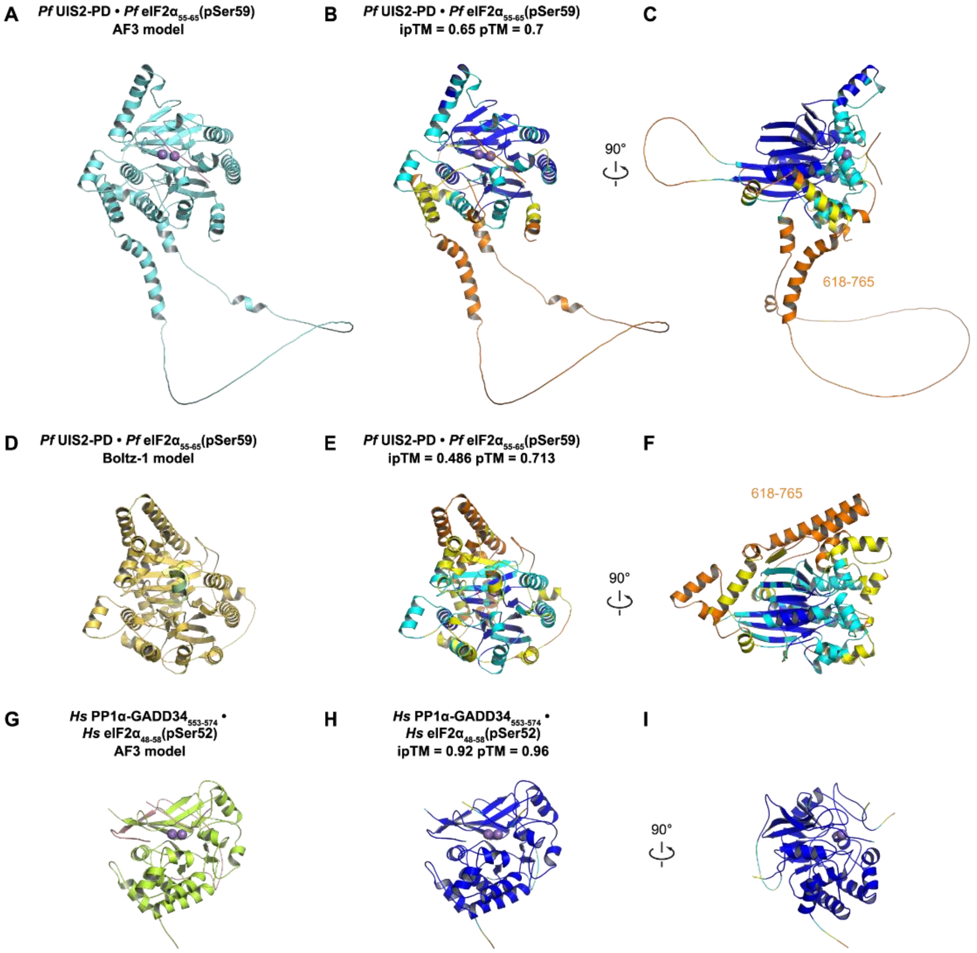


**Supplementary Figure 9. Predicted models of *Pf* UIS2-PD-eIF2α$_{55\text{-}65}$(pSer59) and the human PP1α–eIF2α$_{48\text{-}58}$(pSer52) complexes. (A-C)** Cartoon representations of the *Pf* UIS2-PD (residues 508–1046) bound to the *Pf* eIF2α$_{55\text{-}65}$ peptide phosphorylated at Ser59 (pSer59), predicted by AF3. The interface TM-score (ipTM) and overall TM-score (pTM) are indicated. **(D-F)** Cartoon representations of the *Pf* UIS2-PD (residues 508–1046) bound to the *Pf* eIF2α$_{55\text{-}65}$(pSer59) peptide, predicted by Boltzmann-trained transformer model (Boltz-1). **(G-I)** Cartoon representations of *Hs* PP1 (residues 508–1046) bound to the human eIF2α$_{48\text{-}58}$(pSer52) peptide, predicted by AF3.

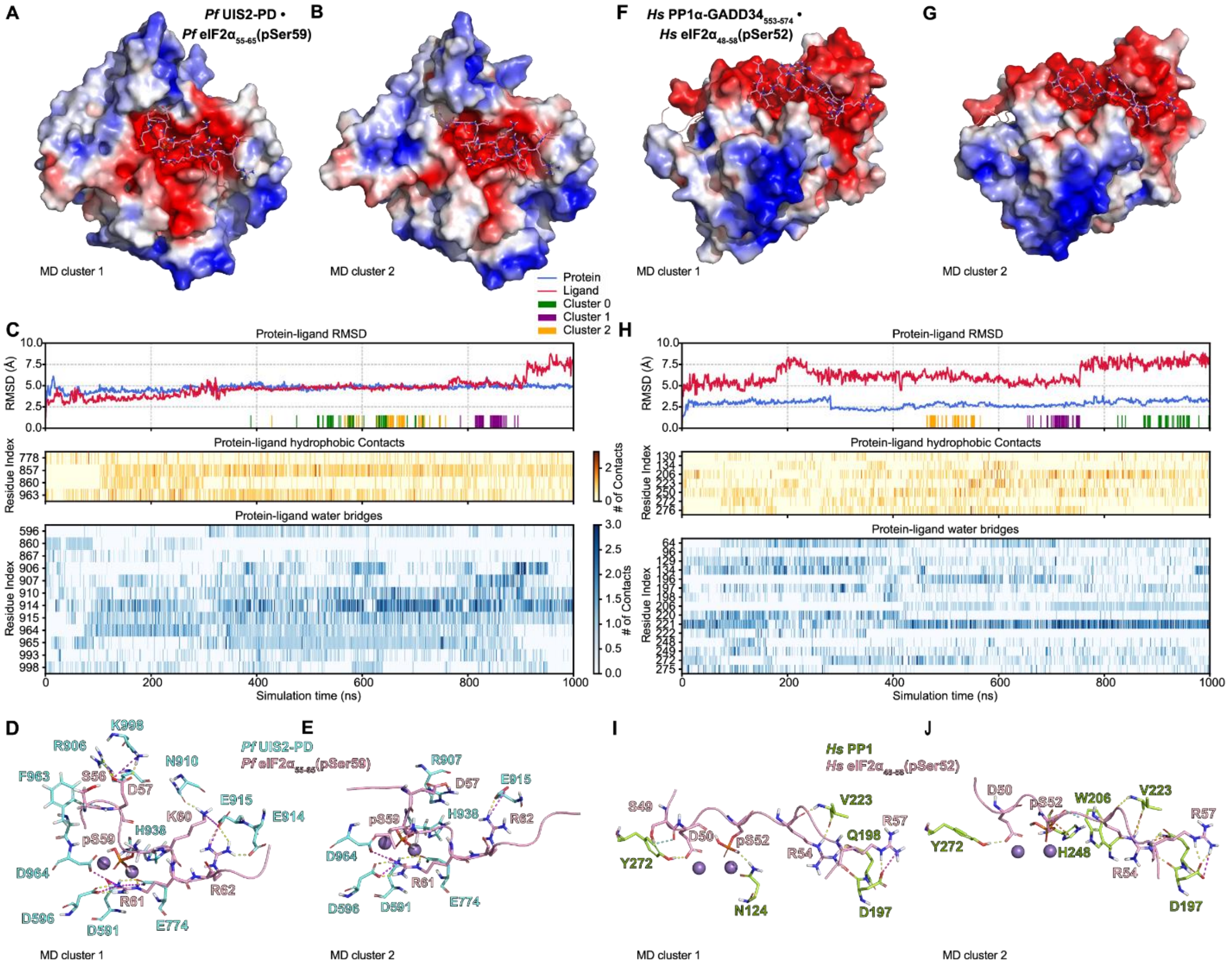


**Supplementary Figure 10. Dynamic binding of phosphorylated eIF2α peptides to *Pf* UIS2 PD and human PP1α, analyzed by molecular dynamics (MD) simulations. (A-B)** Representative structures of clusters 1 and 2 from the *Pf* UIS2-PD–eIF2α(pSer59) complex, sampled during the 1 µs MD trajectory. **(C)** Time traces from the 1 µs MD simulation: backbone RMSD for *Pf* UIS2-PD (blue) and ligand RMSD for eIF2α(pSer59) (red); cluster assignments shown by the color bar; and heatmaps of per-residue hydrophobic contacts and water-mediated interactions. **(D-E)** Cartoon views of key interactions in the cluster 1 and 2 structures. Color code: yellow = hydrogen bond; magenta = salt bridge; cyan = aromatic hydrogen bond. **(F-G)** Representative structures of clusters 1 and 2 from the human PP1α in complex with GADD34$_{553\text{-}574}$ and the human eIF2α$_{48\text{-}58}$ peptide MD trajectory. **(H)** Time traces from the 1 µs MD simulation: backbone RMSD for PP1α (blue) and ligand RMSD for pSer52 peptide (red); cluster assignments indicated by the color bar; and heatmaps of per-residue hydrophobic contacts and water bridges. **(I-J)** Cartoon views of key interactions in the cluster 1 and 2 for the PP1α complex. Color coding follows that in panels **D–E**.

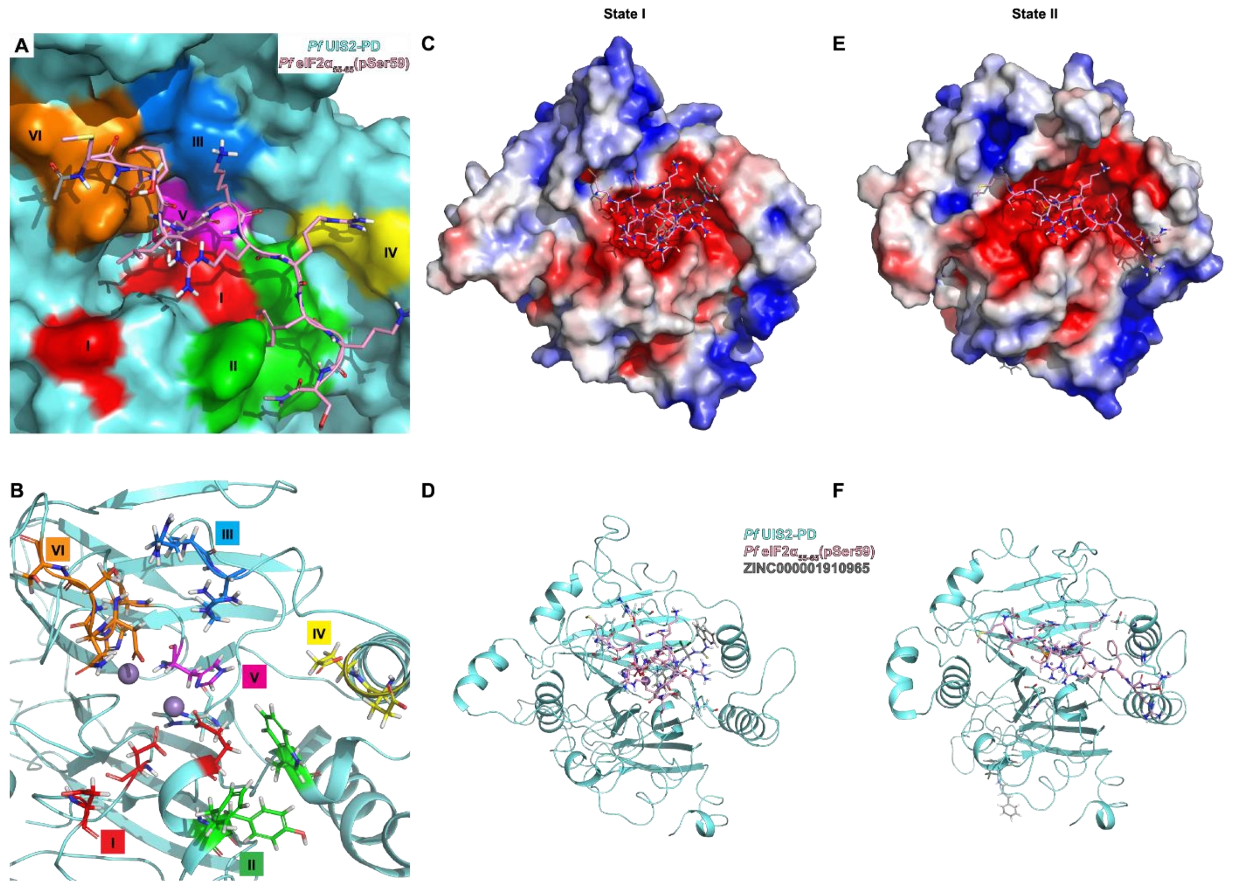


**Supplementary Figure 11. Structural clustering and conformational transitions during Salubrinal and pSer59 loop competition. (A–B)** Surface **(A)** and cartoon **(B)** views of six binding patches. Patches I–VI are defined based on spatial proximity and side-chain chemistry, and collectively mediate UIS2–eIF2α(pSer59) interactions. **(C–D)** Surface **(C)** and cartoon **(D)** snapshots of low-energy State **I**, in which Salubrinal and the pSer59 loop co-occupy the UIS2 binding pocket (see Figs. 5D-E). **(E–F)** Surface **(E)** and cartoon **(F)** Views of low-energy State II, where Salubrinal is dissociated and only the pSer59 peptide remains bound. (see Figs. 5F-G).

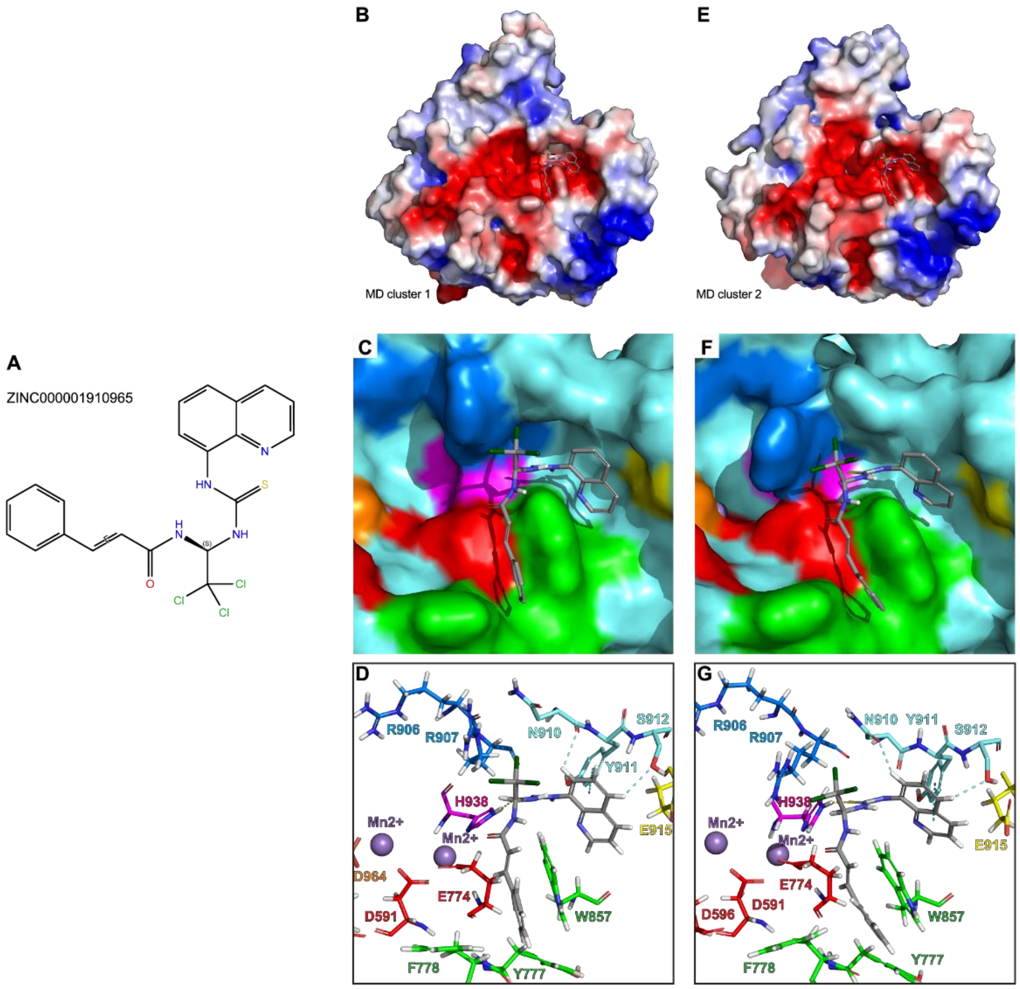


**Supplementary Figure 12. Binding conformations of Salubrinal within the *Pf* UIS2-PD pocket from molecular dynamics (MD) clustering analysis. (A)** 2D chemical structure of Salubrinal. **(B–D)** Representative conformation from Cluster 1 of the MD trajectory (see Fig. 6G). **(B)** Surface representation of the *Pf* UIS2-PD-Salubrinal complex. **(C)** Same view as in (B), with interaction patches color-coded by region (I–VI). **(D)** Cartoon representation with key interacting residues and coordination sites annotated. **(E–G)** Representative conformation from Cluster 2 of the MD trajectory (see Fig. 6G). **(E)** Surface representation of the *Pf* UIS2-PD-Salubrinal complex. **(F)** Same view as in **(E)**, with interaction patches color-coded by region (I–VI). **(G)** Cartoon representation with key interacting residues and coordination sites annotated.
